# Unique longitudinal contributions of sulcal interruptions to reading acquisition in children

**DOI:** 10.1101/2024.07.30.605574

**Authors:** Florence Bouhali, Jessica Dubois, Fumiko Hoeft, Kevin S. Weiner

## Abstract

A growing body of literature indicates strong associations between indentations of the cerebral cortex (i.e., sulci) and individual differences in cognitive performance. Interruptions, or gaps, of sulci (historically known as *pli de passage*) are particularly intriguing as previous work suggests that these interruptions have a causal effect on cognitive development. Here, we tested how the presence and morphology of sulcal interruptions in the left posterior occipitotemporal sulcus (pOTS) longitudinally impact the development of a culturally-acquired skill: reading. Forty-three children were successfully followed from age 5 in kindergarten, at the onset of literacy instruction, to ages 7 and 8 with assessments of cognitive, pre-literacy, and literacy skills, as well as MRI anatomical scans at ages 5 and 8. Crucially, we demonstrate that the presence of a left pOTS gap at 5 years is a robust and relatively specific longitudinal predictor of better future reading skills in children, with large observed benefits on reading behavior ranging from pseudoword decoding to reading comprehension. The effect of left pOTS interruptions on reading acquisition accumulated through time and showed strong predictive power relative to benchmark cognitive and familial predictors of reading ability and disability. Finally, we show that increased local U-fiber white matter connectivity associated with such sulcal interruptions possibly underlie these behavioral benefits, by providing a computational advantage. To our knowledge, this is the first quantitative evidence supporting a potential integrative gray-white matter mechanism underlying the cognitive benefits of macro-anatomical differences in sulcal morphology related to longitudinal improvements in a culturally-acquired skill.

## Introduction

Identifying neuroanatomical features associated with uniquely human cognitive abilities is a central goal in cognitive neuroscience. A growing number of empirical findings identify relationships between the morphology of indentations (known as sulci) in the cerebral cortex and individual differences in cognition (e.g., as reviewed in Cachia et al., 2021; Mangin et al., 2019; Weiner, 2019, 2023). For example, sulcal morphology in various human association cortices that have expanded significantly throughout evolution has been related to individual differences in abstract reasoning, executive function, inhibitory control, memory, face processing, and reading, as well as psychopathology (Amiez et al., 2018; Borst et al., 2014; Cachia et al., 2014; Garrison et al., 2015; Im et al., 2016; Lahutsina et al., 2023; Maboudian et al., 2024; Parker et al., 2023; Tissier et al., 2018; Voorhies et al., 2021; Willbrand et al., 2022; Yao et al., 2022). Moreover, recent research suggests that individual differences in sulcal interruptions are also related to individual differences in cognitive abilities such as numerical processing (Roell et al., 2021; Schwizer Ashkenazi et al., 2024), language and memory (Santacroce et al., 2024), and reading (Borst et al., 2016; Cachia et al., 2018). Sulcal interruptions (hereby referred to as gyral gaps) are a particular case of interconnecting gyri, historically known as pli de passage as coined by Gratiolet (1854), when they emerge on the brain surface. Nevertheless, it is presently unknown whether gyral gaps that are uniquely human are longitudinally (and thus likely causally) related to changes in cognitive abilities, especially in “evolutionarily new” domains such as reading.

For instance, previous work suggests that the maturation of a functional region for reading in ventral temporal cortex (hereafter VTC) when children learn to read, commonly referred to as the visual word form area (VWFA; Cohen et al., 2000; Dehaene et al., 2015; Dehaene & Cohen, 2011), is in part dependent on the pre-existing neural architecture of this region, in terms of both connectivity and cortical morphology (Beelen et al., 2019; Bouhali et al., 2014; Li et al., 2020; Moulton et al., 2019; Saygin et al., 2016; Skeide et al., 2016; Vandermosten et al., 2016). Of note, the human OTS is laterally displaced compared to the OTS in non-human primates (Natu et al., 2021). Additionally, recent studies have linked individual differences in the presence of gyral gaps in this laterally displaced human OTS to individual differences in reading skills, although findings have not been entirely uniform (Borst et al., 2016; Cachia et al., 2018; Del Mauro et al., 2022; Roell et al., 2022).

However, no study has yet explored if this unique neuroanatomical feature in human VTC is related to individual differences in reading skills, as well as other reading precursors (and related cognitive skills such as phonological awareness) as children learn to read - and additionally, if white matter changes may mediate this complex relationship.

Specifically, recent studies that identify relationships between sulcal morphology and cognition propose the novel hypothesis that this relationship reflects underlying differences in white matter architecture. Yet, to our knowledge, this hypothesis has not yet been tested. Sulcal interruptions are particularly interesting to study as they have been linked to a higher density of underlying short range white matter fibers (Bodin et al., 2021) - in which, mechanistically, a higher density of short range white matter fibers could provide a more efficient neuroanatomical infrastructure contributing to individual differences in human cognition compared to a less dense, and therefore, inefficient, neuroanatomical infrastructure. To test these options, here, we investigated whether the presence of left OTS interruptions serve as a longitudinal predictor of reading skills for the first time. We also examine the predictive power of these interruptions relative to benchmark cognitive precursors of reading and assess the role of white matter properties that may mediate this relationship.

Utilizing longitudinal data from 43 children followed from kindergarten at age 5 to age 8 on three measurement occasions, with comprehensive behavioral assessments of literacy, pre-literacy, cognitive skills, and MRI scans, our results provide novel evidence supporting that the interruption of the left posterior OTS (pOTS) is a robust longitudinal predictor of better reading skills, independent from—and strongly predictive relative to —typical pre-literacy skills. This relationship holds and accumulates across time, suggesting that the morphology of the left pOTS is a strong and consistent indicator of reading proficiency. This anatomical marker contributes beyond established behavioral and familial predictors of reading, and this association appears relatively specific to literacy rather than reflecting a general cognitive advantage. Increased local U-fiber structural connectivity associated with left pOTS interruption may underpin the reading benefit of this neural marker. By exploring the neuroanatomical underpinnings of reading, our results highlight the intricate connection between the brain’s evolved structure and its capacity to adapt to culturally invented cognitive demands, such as reading.

## Results

Due to previous findings showing (i) a link between gyral gaps of the occipito-temporal sulcus (OTS) and reading abilities (Borst et al., 2016; Cachia et al., 2018) and (ii) the importance of the neighboring mid-fusiform sulcus (MFS) as a cytoarchitectonic and functional landmark in VTC (Grill-Spector & Weiner, 2014; Weiner, 2019; Weiner et al., 2014), potentially relevant for graphemic orthographic processing (Bouhali et al., 2019), the OTS and MFS were delineated manually in each hemisphere, without awareness of the reading scores. This was done at each time point with available MRI scans (Time 1 at age 5 - T1, and Time 3 at age 8 - T3) on cortical surface reconstructions (in Freesurfer; Dale et al., 1999; Fischl et al., 1999; Reuter et al., 2012; Zaretskaya et al., 2018; see Materials and Methods) obtained by both cross-sectional and longitudinal pipelines (see Figure 1 for data overview and sulcal definitions, as well as Supplementary Figure 1 in Appendix A for a view of sulcal definitions in all participants). Because there were two T_1_-weighted MRI scans per child (at T1 and T3), a within-subject template was created through the Freesurfer longitudinal pipeline to refine brain segmentations at each time point - an approach that increases the precision and discrimination of cortical surface reconstructions (Reuter et al., 2012). Brain-behavior relationships with regard to reading development described were restricted to the left pOTS, and other delineated sulci serve as controls to demonstrate its specificity.

**Figure 1:**
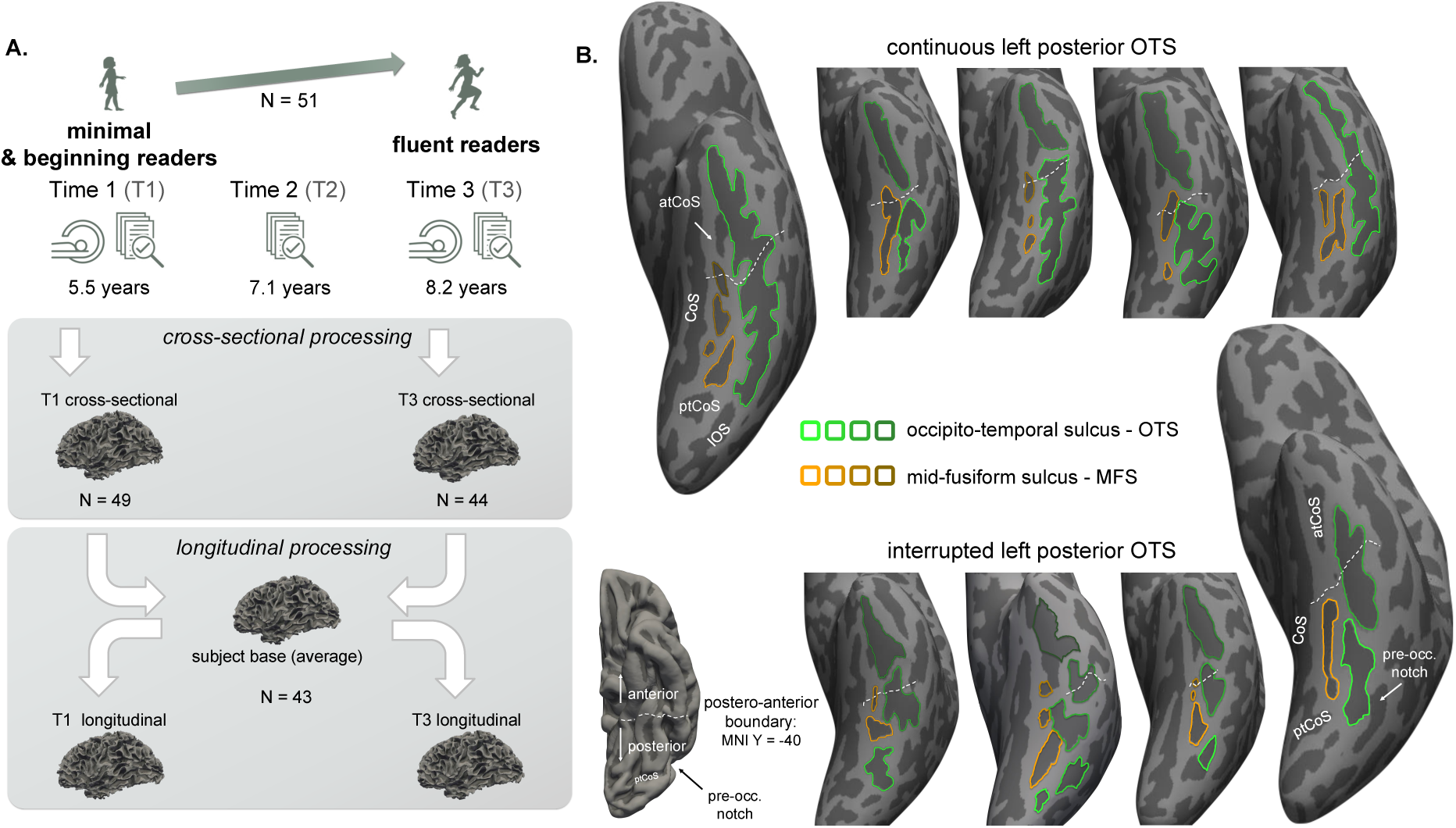
Overview of the experimental design, analytical pipeline, and sulcal definitions. **A.** Children were followed longitudinally from approximately 5.5 years of age (Time 1, T1) to age 8.2 (Time 3, T3) with both behavioral assessments and MRI scans, in addition to an intermediate behavior-only assessment at age 7.1 (Time 2, T2). T_1_-weighted images from T1 and T3 were first processed separately in Freesurfer (cross-sectional processing), and then through the longitudinal pipeline with the creation of a subject template and refinement of T1 and T3 tissue segmentations. **B.** The occipito-temporal sulcus (OTS, green) and mid-fusiform sulcus (MFS, orange) were defined manually, as shown on 9 sample individual inflated left hemispheres (T1 longitudinal processing). Posterior and anterior sections of ventral temporal cortex were defined in each individual based on a boundary (dotted white line) located at MNI Y=-40 (see projection on Freesurfer average (*fsaverage)* surface on the bottom left, which roughly aligned with the anterior tip of the MFS). Left hemispheric views from exemplar children are shown, either with a continuous left posterior OTS on the top row (i.e., no more than one connex OTS component in the space posterior to the white dotted line) or an interrupted left pOTS on the bottom row (two or more sulcal components present in posterior space, with a total Euclidean distance between components ≥1mm). Green and orange gradient colors are respectively used to highlight the different sulcal components of the OTS and MFS. CoS: collateral sulcus; ptCoS/atCoS: posterior/anterior transverse branch of the CoS; IOS: inferior occipital sulcus.

### When present, a gyral gap in the posterior OTS is a reliable longitudinal landmark

The OTS showed a very high incidence of interruption as a whole: approximately 79.3% of children showed at least one interruption in the left OTS, while 74.6% showed interruptions in the right OTS (see Supplementary Figure 2 for interruption statistics for all sulci, time points and processing pipelines). The posterior section of the OTS, whose interruption pattern has been more specifically implicated with reading skills (Cachia et al., 2018), had at least one interruption in the left hemisphere in 56.5% of children, and in 58.3% in the right.

We further evaluated the stability and reliability of left pOTS interruption measurements across T1 and T3. Acceptable internal consistency in measuring the presence of left pOTS interruptions between T1 and T3 was observed when using segmentations from the cross-sectional processing pipeline (T1-T3 Cronbach’s α = .778, estimated 95% confidence interval based on the procedure by Feldt et al. (1987): CI= [.591 - .880]). However, consistency was excellent and significantly improved with the longitudinal processing pipeline (T1-T3 α = .976, CI= [.958 - .976]). Only one child changed left pOTS interruption classification between T1 and T3 under longitudinal processing, due to the measured inter-component distance crossing the predefined 1-mm threshold, consistent with high stability of this macro-anatomical feature. Similarly, other anatomical measures, including sulcal length, sulcal interruption distance, surface area and thickness, also showed higher consistency using Freesurfer’s longitudinal pipeline (see Supplementary Table 1 for the left pOTS). Left pOTS interruptions therefore appeared as a stable anatomical feature in time (at least in the 5- to 8-year-old range), in agreement with the prevailing view that the sulcal morphology of secondary sulci - including interruption patterns - is set in utero (Cachia et al., 2021). Left pOTS interruption was best measured with two T_1_-weighted scans using Freesurfer’s longitudinal (rather than cross-sectional) processing.

Because our interest lies in how sulcal morphology prospectively predicts the emergence of reading skills over time, we hereafter report the association between reading skills and left pOTS interruption measured through the longitudinal pipeline at Time 1 (T1). We report results from all 43 children who could be processed through this longitudinal pipeline (see participants’ characteristics in Table 1). However, similar results are observed regardless of type of processing (longitudinal vs. cross-sectional) and time point used for the analyses (see Appendix A - Figure S3A).

**Table 1:** Sample demographics and main neuropsychological scores at the three time points T1-3. (median and inter-quartile range are indicated), for the N=43 children with longitudinal MRI processing reported in the main text.

|  | Full longitudinal sample<br>(N=43) | Initial minimal readers<br>(N=24, subset) |
| --- | --- | --- |
| Gender | 18 Females / 25 Males | 8 Females / 16 Males |
| Handedness | 38 Right / 5 Left | 23 Right / 1 Left |
| Family history of dyslexia | 18 + / 25 - | 8 + / 16 - |
| Age T1 | 5.49 years [5.31 - 5.86] | 5.47 [5.19 - 5.71] |
| Age T2 | 7.13 years [6.87 - 7.50] | 7.10 [6.79 - 7.46] |
| Age T3 | 8.18 years [8.00 - 8.57] | 8.20 [8.04 - 8.54] |
| TOWRE reading scores T1 |  |  |
| SWE (raw score) | 3.0 [0.0 - 9.0] | 0.0 [0.0 - 2.3] |
| PDE (raw score) | 2.0 [0.0 - 5.0] | 0.0 [0.0 - 1.0] |
| TOWRE reading scores T2 |  |  |
| SWE (raw score) | 49.0 [38.0 - 58.5] | 44.0 [31.0 - 53.3] |
| PDE (raw score) | 18.0 [13.0 - 24.5] | 18.5 [13.0 - 24.0] |
| SWE (standard score) | 109.0 [103.0 - 120.0] | 107.5 [98.8 - 115.5] |
| PDE (standard score) | 104.0 [99.0 - 112.0] | 105.0 [99.0 - 111.5] |
| TOWRE composite (standard score) | 108.0 [101.0 - 118.0] | 108.0 [100.5 - 116.2] |
| TOWRE reading scores T3 |  |  |
| SWE (raw score) | 61.0 [55 - 67.5] | 58.5 [50.5 - 69.0] |
| PDE (raw score) | 24.0 [20 - 35.5] | 24.0 [17.0 - 36.5] |
| SWE (standard score) | 110.0 [103.5 - 118.0] | 106.5 [101.8 - 118.0] |
| PDE (standard score) | 104.8 [98.0 - 115.0] | 100.5 [96.0 - 116.2] |
| TOWRE composite (standard score) | 108.9 [101.0 - 116.0] | 104.5 [97.5 - 119.0] |
| Rapid Automatized Naming<br>(standard score, average of color and object<br>naming) |  |  |
| T1 | 98.0 [89.5 - 107.5] | 97.0 [88.9 - 103.1] |
| T2 | 97.0 [88.3 - 107.8] | 97.3 [84.8 - 104.1] |
| T3 | 97.0 [84.5 - 105.3] | 95.8 [84.5 - 104.0] |
| Phonological awareness (CTOPP)<br>(standard composite score of elision &<br>blending subtests) |  |  |
| T1 | 112.0 [104.5 - 118.0] | 106.0 [100.0 - 112.0] |
| T2 | 118.0 [107.5 - 131.5] | 118.0 [111.2 - 133.8] |
| T3 | 119.8 [109.0 - 127.0] | 118.0 [108.2 - 127.0] |
| Letter Knowledge T1 (WMRT)<br>(standard score) | 110.0 [103.5 - 114.5] | 107.0 [102.8 - 110.2] |
| Performance IQ (WJC Concept Formation,<br>standard score) |  |  |
| T1 | 119.0 [106.0 - 127.0] | 113.5 [103.0 - 123.5] |
| T3 | 117.0 [109.0 - 123.0] | 117.0 [107.5 - 121.0] |
| Socio-economic status |  |  |
| Average parental years of education | 17 years [16 - 18] | 16 years [15.5 - 17] |
| Yearly household income | 140k\$ [88.5 - 200] (*N=39) | 150k\$ [92.5 - 200] (*N=23) |

### Left pOTS interruption is prospectively associated with better reading skills

The presence of interruptions in the left pOTS at T1, measured with the longitudinal processing pipeline, was significantly associated with reading skills measured at T3. Compared with children with a continuous pOTS, children with an interrupted left pOTS at T1 showed higher reading TOWRE composite standard scores at T3, while controlling for demographic variables and total brain volume (see Figure 2A; F-test comparing nested models predicting TOWRE scores by age at T3, handedness and gender only, or by adding the residuals of a logistic regression explaining left pOTS interruption by total brain volume, age at T1, gender and handedness: F(1, 38) = 11.34, p = 1.7*10^-3^, ΔR^2^ = 20.5%). This corresponded to a large effect size (Cohen’s d on reading residuals accounting for demographic variables: d = 0.957).

**Figure 2:**
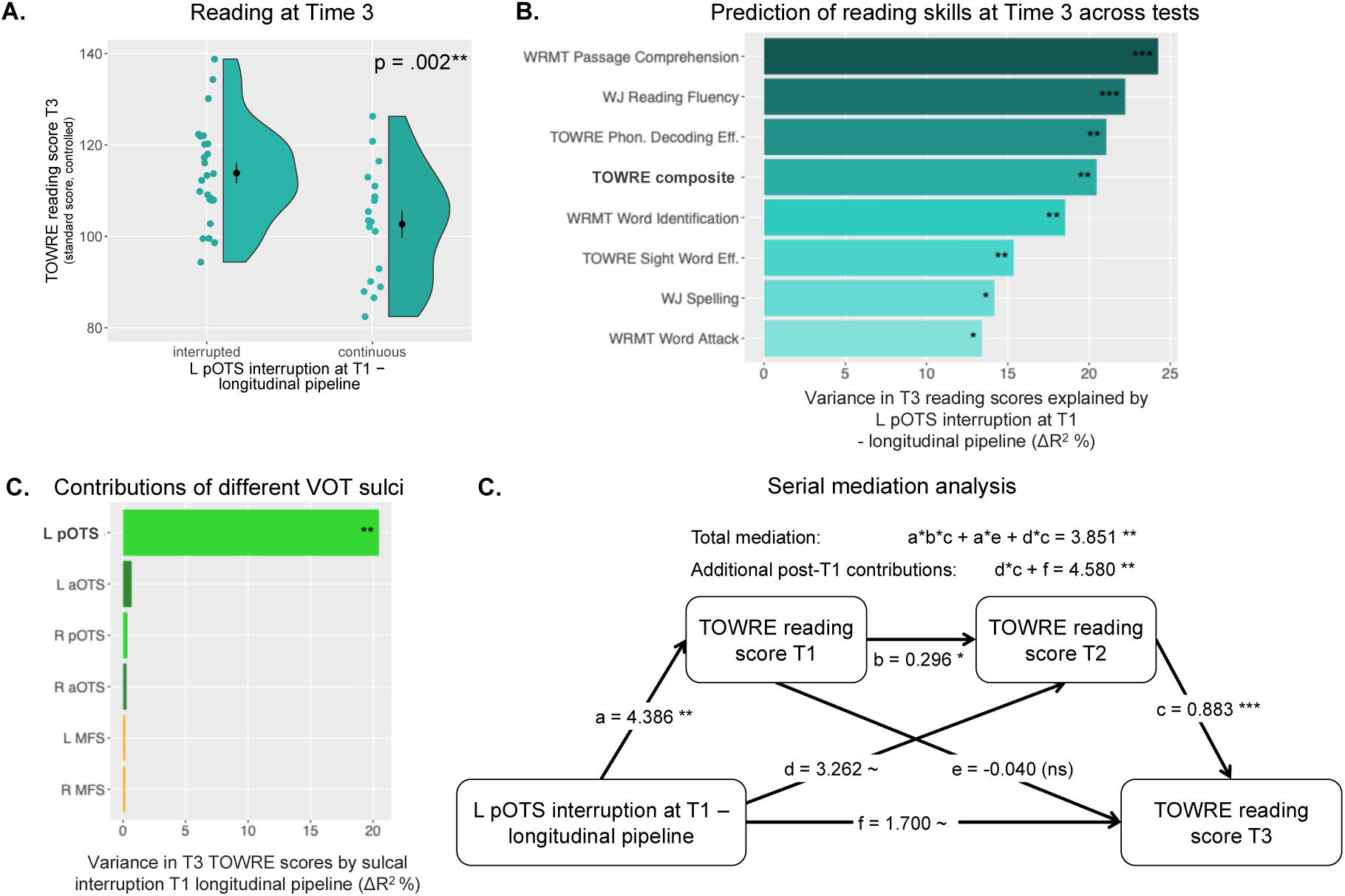
Cumulative impact of left pOTS interruption on reading scores longitudinally. **A.** TOWRE composite standard scores at T3 are significantly higher in children with an interrupted than continuous left pOTS (assessed at T1 with longitudinal processing), after removing residual variance due to age, gender and handedness. Black dots represent mean scores in each group, ± standard error of the mean (s.e.m.). **B. Prediction of reading and spelling scores across tests:** percentage of variance in different reading and spelling standardized tests explained by left pOTS interruption (ΔR^2^ in models controlling for demographic variables and total brain volume). Left pOTS interruption at T1 was significantly associated with all reading scores at T3, ranging from timed and untimed measures of single word and pseudo-word reading, as well as sentence and passage comprehension or spelling. The bolded TOWRE composite score corresponds to the effect depicted in panel a. **C. Percentages of variance in TOWRE T3 reading scores** explained by interruption by different sulci in VTC (posterior and anterior OTS (pOTS and aOTS, respectively) and MFS, in each hemisphere). **D. Serial mediation model demonstrating the cumulative impact of left pOTS interruption on TOWRE reading scores over time** from T1 to T2 to T3. Coefficients are standardized. Reading scores are controlled for age at testing, gender, and handedness, as well as left pOTS interruption by these demographic variables and total brain volume (***: p<0.001; **: p<.01, *: p<.05).

The association between left pOTS interruption and reading skills was robust to different measurements of reading skills and sulcal interruption in four ways. First, left pOTS interruption (measured at T1 with longitudinal processing) was predictive of reading skills as measured by all reading tests at T3, with a range of ΔR^2^ = 13.4% to 24.3% of variance explained on a variety of reading tests, ranging from accuracy and speed/fluency reading aloud single words and pseudo-words, to silent sentence reading and passage comprehension, as well as spelling scores (see Figure 2B). All of these associations were statistically significant (p < .012). Interruption of the left pOTS was most strongly associated with passage comprehension and sentence reading scores.

Second, associations between sulcal interruption and reading skills were specific to the left pOTS within VTC. No significant associations were observed for anterior OTS (aOTS) or MFS in either hemispheres, or for the right pOTS (see Figure 2C, all p-values > 0.58, all ΔR^2^ < 0.7%).

Third, left pOTS interruption predicted T3 TOWRE composite scores regardless of whether it was measured through the longitudinal or cross-sectional processing pipeline of Freesurfer, at T1 or T3 (see Supplementary Figure 3A; T3 longitudinal: F(1, 38) = 9.57, p = 3.7*10^-3^, ΔR^2^ = 17.9%; T1 cross-sectional: F(1, 40) = 10.96, p = 2.0*10^-3^, ΔR^2^ = 19.6%; T3 cross-sectional: F(1, 39) = 3.20, p = .08, ΔR^2^ = 6.62%), although the contribution was marginal at T3 with the cross-sectional pipeline. This importantly implies that left pOTS morphology at T1 was still a significant longitudinal predictor of T3 reading behavior when T3 brain images were not used to improve T1 anatomical segmentations, i.e., with T1 cross-sectional processing alone, despite noisier tissue segmentations with cross-sectional processing (see also Reuter et al., 2012).

Fourth, we tested whether interruption distance, as a continuous measure of sulcal interruption rather than the binary presence or absence of a gyral gap, was more predictive of reading skills. The total interruption distance between segments of the left pOTS was marginally predictive of future reading skills (F(1, 38) = 3.54, p = .067, ΔR^2^ = 7.6%, see Supplementary Figure 3B). However, two additional analyses suggest that this association is likely due to trivial correlations between interruption distance and the binary presence of an interruption: (i) a similar analysis restricted to the subsample of children with interrupted left pOTS showed no association between reading and left pOTS total interruption distance (F(1, 21) < 0.001, ΔR^2^ = 0.3%, p = .98); and (ii) interruption distance does not explain additional variance over and above the binary presence of an interruption across all children (F(1, 37) = 0.04, p = .85), while binary interruption significantly explains more variance in reading skills over and above interruption distance (F(1, 37) = 6.99, p = .012).

We further tested other anatomical features of the left pOTS at T1 and found that no other feature considered predicted reading skills, including total sulcal length, sulcal depth, depth of the sulcal pit, mean thickness, and total surface area (see Supplementary Figure 3B, all p-values > .26). In sum, the prospective relationship between left pOTS morphology at T1 and reading skills at T3 was specific to the binary presence of a gyral gap.

### The effect of left pOTS interruption on reading accumulates over time

We next tested whether the effect of left pOTS interruption on reading scores observed at T3 was present at earlier time points and how it evolved over time. Compared with children with a continuous left pOTS, children with interrupted left pOTS also had higher TOWRE composite reading scores at T2, at around 7.1 years-old, and already at T1, at approximately 5.5 years-old (see Supplementary Figure 4B and C; T2 TOWRE composite scores: F(1, 38) = 9.43, p = 3.9*10^-3^, ΔR^2^ = 14.3%, Cohen’s d = 0.872; population-normalized T1 TOWRE composite scores, in the absence of norms in this age-range: F(1, 38) = 7.63, p = 8.8*10^-3^, ΔR^2^ = 16.0%, Cohen’s d = 0.919).

However, differences in emerging reading skills at T1 did not alone drive the later differences observed at T2 and T3. To test this, we conducted a supplementary analysis in the subset of 24 children with the lowest reading scores at T1 (defined as those with a TOWRE T1 population-derived composite score equivalent below 95). Hereafter, we refer to these children as *initial minimal readers*: 79% could not decode a single syllable or pseudoword in 45 s, with a maximum of four pseudowords decoded, and 63% could not recognize a single frequent sight word, with a maximum of seven words recognized. Although these children showed no difference in T1 reading scores as a function of whether their left pOTS was interrupted or continuous (F(1, 19) = 0.21, p =.66, ΔR^2^ = 0.82%, Cohen’s d = 0.034), there were growing statistically significant differences at T2 (F(1, 19) = 5.20, p =.034, ΔR^2^ =16.91%, Cohen’s d = 0.965) and at T3 (F(1, 19) = 6.93, p =.016, ΔR^2^ = 26.15%, Cohen’s d = 1.107; see Supplementary Figure 4D-F).

A serial mediation analysis confirmed that the beneficial effect of left pOTS interruption on reading acquisition accumulated over time, from T1 to T2 and T3, and did not merely result from differences already present at T1 (see Figure 2C). Residuals of T1, T2 and T3 TOWRE reading scores, controlled for age, gender and handedness, and residuals of a logistic regression of left pOTS interruption, accounting for demographic variables and total brain volume, were entered into a serial mediation model. T1 and T2 reading scores were significant serial mediators of the effect of left pOTS interruption on T3 reading (total serial mediation a*b*c+a*e+d*c: estimated coefficient ꞵ = 3.851, standard error SE = 1.450, p = .008). Notably, left pOTS interruption at T1 predicted T1 TOWRE score (path a: ꞵ= 4.386, SE=1.560, p = .004), T1 TOWRE predicted T2 TOWRE (b: ꞵ = 0.296, SE = 0.149, p = .049) and T2 TOWRE predicted in turn T3 TOWRE (c: ꞵ = 0.883, SE = 0.104, p < .001, see Figure 2C). Importantly, there were also significant contributions of left pOTS interruption onto T3 reading on top of those mediated through T1, with large cumulative effects through T2 and directly onto T3 (d*c+f: ꞵ = 4.580, SE = 1.757, p = .009), while indirect effect through T1 TOWRE scores alone were non-significant (a*b*c+a*e: ꞵ = 0.971, SE = 0.839, p = .25). Left pOTS interruption had marginal direct effects both onto T2 TOWRE scores (d: ꞵ = 3.262, SE = 1.673, p = .053) and onto T3 TOWRE (f: ꞵ = 1.700, SE = 0.983, p < .09). Interestingly, the magnitude of direct contributions of left pOTS interruption onto reading scores seemed to decrease over time (a: ꞵ = 4.386, d: ꞵ = 3.262, f: ꞵ = 1.700). Overall, left pOTS interruption therefore appeared as a continued constraint on reading acquisition between 5 and 8 years of age.

### Left pOTS interruption makes a strong longitudinal contribution relative to established predictors of reading

Having observed that left pOTS interruption before or at the onset of literacy acquisition is a strong predictor of future reading skills, we next asked how predictive this neuroanatomical feature was relative to benchmark cognitive predictors of reading skills, and how these variables related to one another. Specifically, we asked whether left pOTS interruption was a predictor of reading independent from known pre-literacy skills, or whether an association with a specific pre-literacy skill before learning to read may trivially explain its predictive power. We considered constructs shown to predict reading acquisition or risk of dyslexia longitudinally, as measured by classical neuropsychological tests: rapid naming (of colors and objects, as categories independent from early reading skills), phonological awareness, performance IQ, and receptive vocabulary. Letter knowledge and TOWRE scores at T1 were also considered, as emerging reading skills. Family history of dyslexia was also taken into consideration, and indexed by mother scores in the Adult Reading History Questionnaire (ARHQ, Lefly & Pennington, 2000, see Methods for reasons for selecting mother ARHQ).

Relative to all these predictors, left pOTS interruption was the strongest longitudinal predictor of reading, with its 20.5% of variance in T3 TOWRE explained after controlling for demographic variables (ΔR^2^ = 20.5%, F(1, 38) = 11.34, p = 1.7*10^-3^), followed by letter knowledge with 19.6% of variance (letter identification, F(1, 38) = 10.75, p = 2.2*10^-3^) and phonological awareness with 16.2% (CTOPP, F(1, 38) = 8.51, p = 5.9*10^-3^; see Figure 3A). Rapid naming (RAN objects and colors), early reading scores (TOWRE T1), family history of dyslexia (mother ARHQ) and receptive vocabulary (PPVT) were also significant but weaker predictors of T3 reading skills, with separate respective contributions to T3 TOWRE scores of 13.6%, 11.5%, 11.2% and 9.5% of variance (p ∈ [0.01, 0.04]). In the subsample of children with minimal reading skills at T1, left pOTS interruption was the second strongest longitudinal predictor of reading explaining 26.1% of variance in T3 reading (F(1,19) = 6.93, p = .02), close after rapid naming (ΔR^2^ = 28.0%, F(1, 19) = 7.61, p = .01, see Figure 3D). Dyslexia family history was the third and only other significant predictor in this subsample (ΔR^2^ = 21.3%, F(1, 19) = 5.29, p = .03; all other ps > .09).

**Figure 3:**
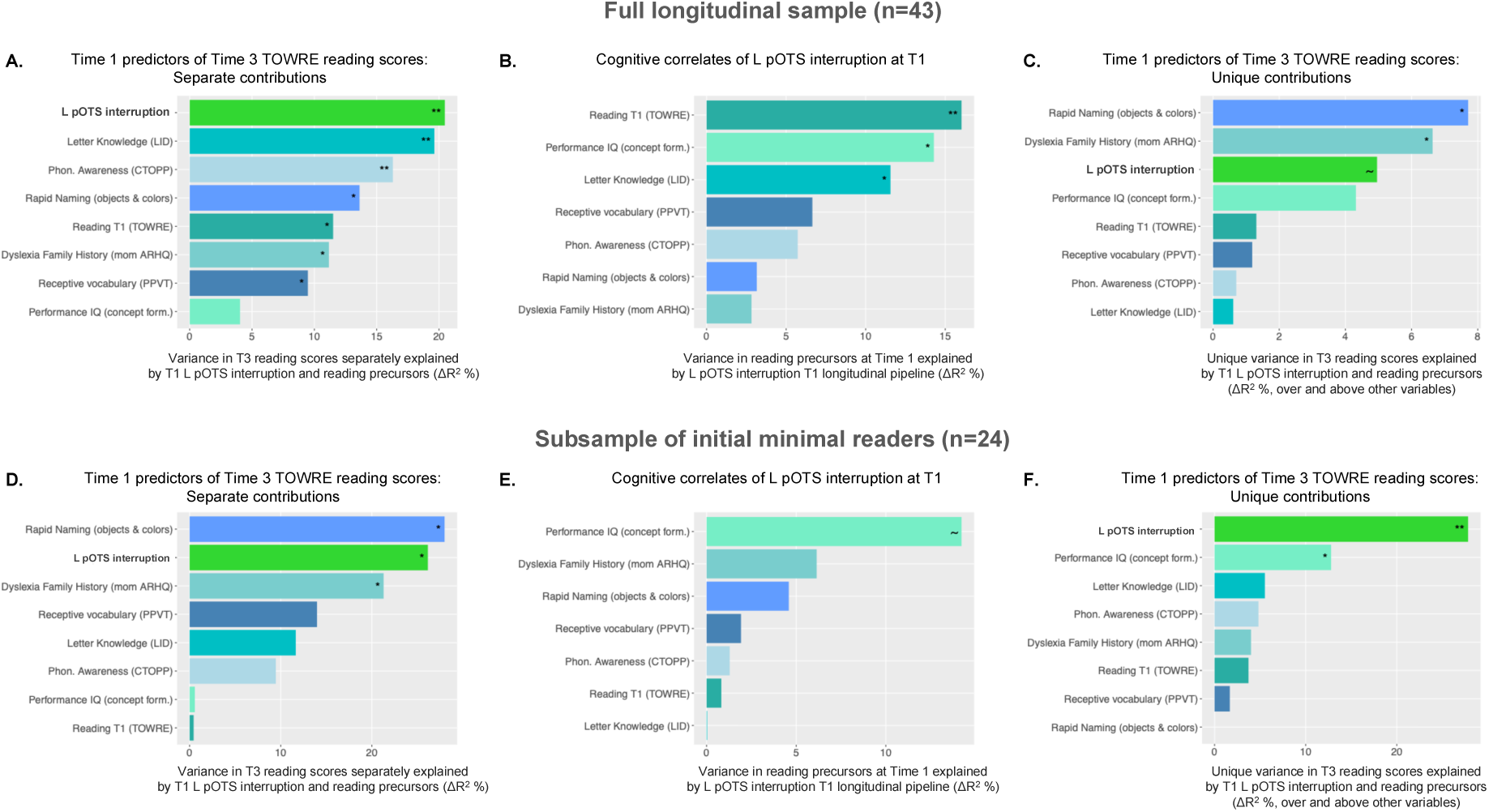
Longitudinal contributions of left pOTS interruption to reading relative to cognitive precursors of reading, as well as their associations, in the full longitudinal sample (A-C, top row) and in the sub-sample of initial minimal readers (D-F, bottom row), respectively. A & D. Variance in TOWRE reading scores at Time 3 explained by left pOTS interruption and classical predictors of reading at Time 1 separately, over and above demographic variables, ranked by decreasing contributions (ΔR^2^ in percentage, relative to baseline reference models with only demographic variables accounting respectively for a total R^2^ of 11.0 and 2.2% of variance in T3 reading). B & E. Associations between left pOTS interruption and predictors of reading at Time 1: percentage of variance (ΔR^2^) in left pOTS interruption explained by each pre-literacy skill separately, over and above demographic variables and total brain volume. C & F. Unique variance in TOWRE T3 reading scores explained by left pOTS interruption and classical predictors of reading at Time 1, over and above other variables of interest and demographic variables, ranked by decreasing contributions (ΔR^2^ in percentage). For reference, the full models including all considered predictors accounted respectively for a total R^2^ of 58.9 and 75.8% of variance in T3 reading. (Stars on each bar denote significant contributions as assessed by nested model comparison using an F-test. **: p<0.01, *: p<.05)

Concerning the extent to which left pOTS interruption is related at T1 to other longitudinal predictors of reading, we observed significant associations between sulcal interruption and early reading skills as previously reported, with left pOTS interruption accounting for ΔR^2^ = 16.0% of variance in T1 TOWRE scores already above demographic variables and total brain volume (F(1, 38) = 7.63, p = 8.8*10^-3^; see Figure 3B), and for 11.6% of letter knowledge, which can be considered an early literacy measure (F(1, 38) = 6.09, p = .018). We also observed significant sulcal contributions for explaining performance IQ (concept formation: ΔR^2^ = 14.3%, F(1, 38) = 6.78, p = .013), but not for other variables (p>.08). The same analysis repeated in the subsample of 24 initial minimal readers at T1, i.e. removing confounds with early reading skills, still showed an association between left pOTS interruption and performance IQ (ΔR^2^ = 14.2%, F(1, 19) = 3.42, p = .0797, of marginal significance likely due to the reduced sample size given the effect size similar to that observed in the full sample), but no significant association with other pre-literacy skills (see Figure 3D). Thus, we observed an association between left pOTS interruption and performance IQ at T1, apparently independent from early reading skills, while the association with letter knowledge and TOWRE scores at T1 may only reflect early effects of left pOTS interruption on emerging reading skills observable in the full sample only.

These significant associations between left pOTS interruption, performance IQ and pre-literacy skills (Figure 3B and D), as well as potential associations between pre-literacy skills themselves (cf. correlation matrices in Supplementary Figure 5D and H), make it challenging to fairly assess the separate unique contributions of each of these variables to the longitudinal prediction of reading skills. For instance, because left pOTS interruption explains a significant amount of variance at T1 in letter knowledge, early TOWRE scores and performance IQ, some of its true direct effect on future T3 reading skills is already captured in a baseline model including these cognitive predictors only, even before left pOTS interruption is explicitly entered as a predictor. Yet, even when controlling in a such a stringent way for all pre-literacy skills, early reading abilities, family history of dyslexia and their shared variance with left pOTS interruption in such nested model comparison, left pOTS interruption still had a marginal additional contribution to predicting T3 TOWRE reading scores (ΔR^2^ = 5.0%, F(1, 31) = 3.74, p = .062) in spite of this important collinearity. Rapid naming and family history were the only two other predictors that outperformed left pOTS interruption in terms of unique contributions to T3 reading (ΔR^2^ = 7.7% and 6.6% for RAN and mother ARHQ respectively, see Figure 3C), possibly because these predictors were among the least confounded with others (see Figure S5D for correlations among predictors) and thus among the least affected by collinearity. Letter knowledge, in contrast, had the smallest, non-significant unique contributions to T3 reading (ΔR^2^ = 0.61%, p = .50). In order to reduce the collinearity confounds resulting from the associations between pOTS interruption and T1 early reading skills, we also reran the same analysis in the subsample of initial minimal readers. Left pOTS interruption in this subsample significantly contributed to predicting future T3 reading over and above all pre-literacy skills and family history (ΔR^2^ = 27.8%, F(1, 12) = 13.80, p = .003), making it the first independent longitudinal contributor of reading (see Figure 3F). The second largest longitudinal predictor in terms of unique contributions was performance IQ (ΔR^2^ = 12.8%, F(1, 12) = 6.34, p = .027). No other predictor had a significant unique contribution to the longitudinal prediction of T3 reading in the subset initial minimal readers, including rapid naming even though its separate contribution was larger than that of pOTS.

In summary, after taking into account its collinearity with early T1 literacy and reducing it in the subsample of minimal initial readers, left pOTS interruption at around 5 years of age emerged as a particularly strong independent longitudinal contributor to reading skills at around 8 years, with predictive performance comparable to or greater than that of classical cognitive and familial predictors of reading ability and disability (dyslexia) that are more proximal to reading behavior.

### Left pOTS interruption has a relatively specific cognitive impact on reading abilities

The incidental observation of a significant association between left pOTS interruption and performance IQ at T1, although smaller in magnitude and not sufficient to account for the association with literacy skills, raises the possibility that left pOTS interruption may confer a broader cognitive advantage rather than a specific benefit for reading acquisition. We therefore conducted a supplementary analysis to examine the relative cognitive specificity versus domain generality of this association. Of note, reading and general cognitive skills are strongly correlated and mutually influence each other across development (Ferrer et al., 2007; Peng & Kievit, 2020; Ünal et al., 2023), making this question of cognitive specificity inherently difficult to address.

We gathered all cognitive and linguistic scores obtained at T3, including measures of mathematics, general cognition, executive functioning, language, and reading (see Table S2 for a full list and description of the included subtests), and tested how each measure was associated with left pOTS interruption, residualized for demographic variables and total brain volume. Left pOTS interruption showed strong and significant associations with several measures of language, reading, mathematics, and general cognitive ability (see Supplementary Figure 6). As expected, we also observed strong and significant correlations among the behavioral measures themselves.

However, the strongest correlations with left pOTS interruption residuals were observed for reading scores. Moreover, hierarchical clustering grouped left pOTS interruption with a sub-cluster that included most reading assessments - spanning phonological decoding, word reading, and sentence reading fluency -, as well as spelling and more surprisingly an expressive language index. This reading-related cluster was most closely related to another cluster comprising measures of written and oral language comprehension (i.e., receptive language), followed by a cluster of phonological awareness. In contrast, measures of mathematics, general cognitive ability, backward working memory and rapid naming formed a separate cluster. Thus, although left pOTS interruption is associated with several cognitive, linguistic, and academic measures, its pattern of associations is most closely aligned with children’s reading outcomes, supporting a relatively specific cognitive advantage for reading.

### Why are interruptions beneficial? Left pOTS interruption and local white-matter connectivity advantages

The underlying mechanisms contributing to why the interruption in the cortical folds around the left pOTS region may so strongly and reliably favor adequate reading acquisition remains unknown. Previous work investigating the relationship between sulcal morphology and behavior have proposed a mediating role of the underlying white matter (Amiez et al., 2018; Borst et al., 2016; Cachia et al., 2018; Garrison et al., 2015; Miller et al., 2021; Voorhies et al., 2021), although these potential mechanisms have not been explicitly explored.

In the case of sulcal interruptions, two explanations stand out as potential mechanisms underlying the behavioral benefits for reading of left pOTS interruptions. First, given that white-matter fibers in general, and especially long-distance projection fibers, tend to terminate more in gyri than sulci (Dannhoff et al., 2023; Ge et al., 2018), having an extra gyral crown, or superficial *pli de passage* (Mangin et al., 2019), in the left pOTS may allow to pack more connections, especially long-distance connections through e.g., potentially a combination of fibers from the (posterior) arcuate fasciculus, vertical occipital fasciculus and inferior longitudinal fasciculus (Bouhali et al., 2014; Grotheer et al., 2021; Moulton et al., 2019; Weiner et al., 2017; Yeatman et al., 2012). Such increased long-distance connections may support a more efficient reading network. Alternatively, the presence of a *pli de passage* in the left pOTS may allow for better local white-matter connectivity through short-range U-shaped fibers traveling under the cortical gray matter. Indeed, Bodin and colleagues (2021) showed that, in the superior temporal sulcus (STS), the presence of *plis de passage* was associated with a higher density of U-shaped fibers connecting adjacent gyri. Superficial *plis de passage*, i.e., complete sulcal interruptions with a gyral gap visible on the cortical surface, showed an even higher local connectivity advantage than deep *plis de passage*, the complete gyral gaps or partial interruptions with sulcal pinching which are buried within the depth of the sulcus. (Please note that in our study, we focus on superficial gyral gaps visible on the cortical surface only, and do not classify deep *plis de passage*.)

In order to test these hypotheses, we leveraged diffusion-weighted data available from Time 3 (n=29 children with data of sufficient quality, see Methods; too few children having usable diffusion data at T1). Gyral gaps between left pOTS sulcal components were manually delineated at the level of left pOTS interruptions (see Figure 4A, and Supplementary Figure 7) and we assessed white-matter properties and connectivity around left pOTS gaps vs. left pOTS sulcal components (what we refer to as pOTS proper) in the same individuals (two leftmost violin plots in Figure 4B, top) and then also between individuals (individuals with uninterrupted pOTS and pOTS proper in individuals with an interrupted pOTS; right two violin plots in Figure 4B, top).

**Figure 4:**
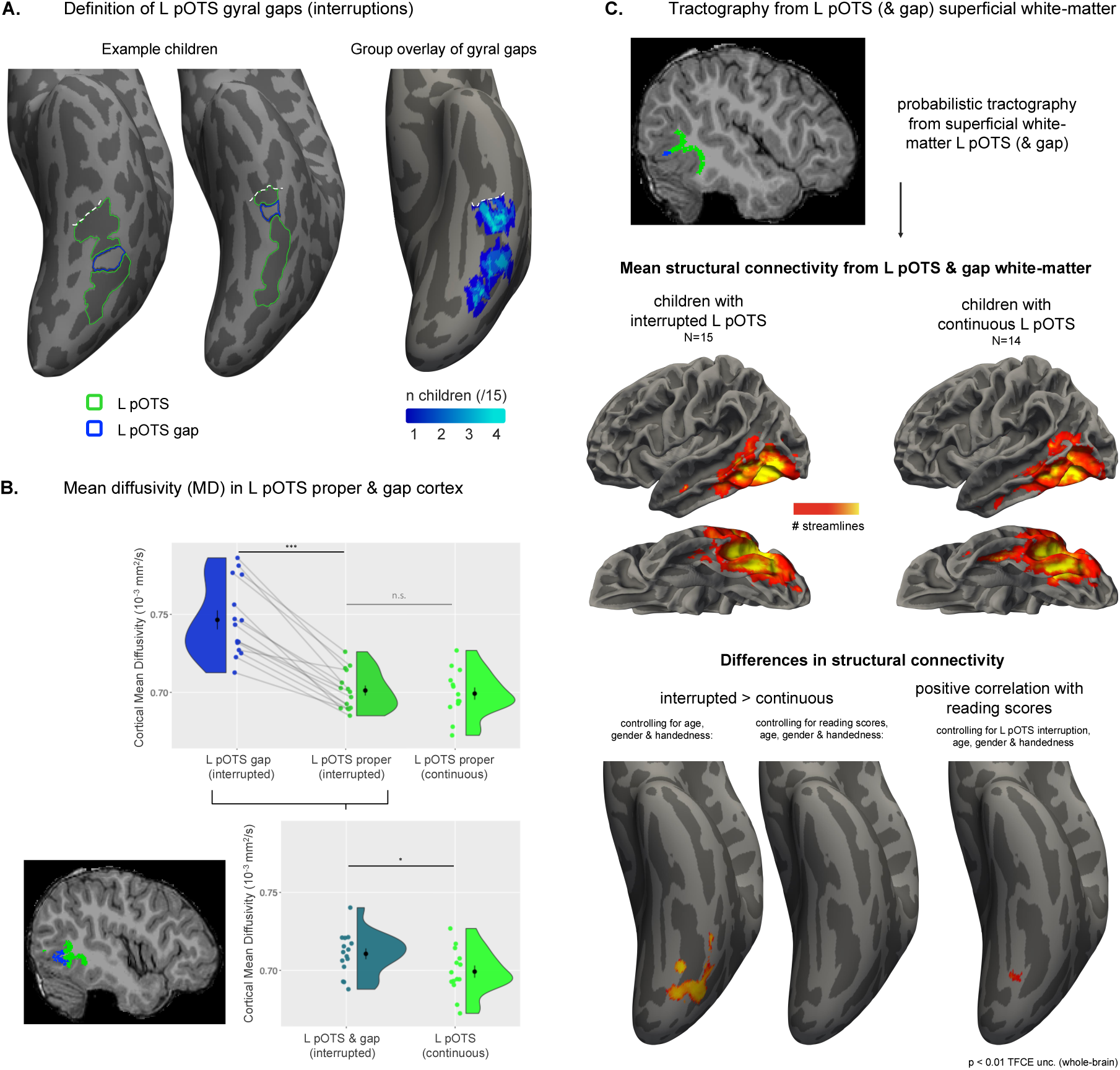
White-matter correlates of left pOTS interruptions. **A.** Gyral gaps in the left pOTS were delineated manually (in blue) and contrasted to left pOTS proper (green). Two exemplar left hemispheres are shown in children with a left pOTS interruption. The group-level map shows overlays of left pOTS gaps (projected onto the *fsaverage* surface) of 15 children with interrupted left pOTS, out of 29 with usable diffusion data. **B.** Average MD in the cortex of the left pOTS proper and gyral gaps. Top row (from left to right): difference in cortical MD in the left pOTS gap (blue) and left pOTS proper (dark green) in children with an interrupted sulcus, and in left pOTS in children with continuous sulci for comparison (light green). Bottom row, left: example cortical regions in which MD was average in a child with a sulcal interruption. Bottom row, right: MD comparison between children with interrupted and continuous sulci over the left pOTS cortex, including left pOTS gap if applicable. **C.** Probabilistic tractography from superficial white-matter seeds above the left pOTS cortex and possible left pOTS gap. From top to bottom: example seed in a participant (green: left pOTS proper, blue: left pOTS gap); mean whole-brain connectivity in children with interrupted (n=15) and continuous left pOTS (n=14); statistical differences in connectivity (from left to right): between the two groups not taking into account reading scores, between the two groups when modeling reading scores (no association observed), between connectivity and T3 TOWRE reading scores (p<0.01 TFCE uncorrected threshold).

According to hypothesis (i), left pOTS gaps should show higher myelination, both in the cortex and the white-matter in its vicinity, than surrounding left pOTS components in the same individuals. Diffusion tensor models were therefore fitted to derive indices related to myelin content. Average Mean Diffusivity (MD) was first estimated in the cortical ribbon as the most robust DTI index reflective of myelination and neurite properties in the cortex (Fukutomi et al., 2019). In children with interruptions (n=15), MD in left pOTS gaps was higher than in the cortex of the left pOTS proper (paired two-sample T-test t(14) = 9.74, p = 1.3*10^-7^ ; see Figure 4B, top row), while MD in the left pOTS proper of these children was similar to MD in the continuous left pOTS of the remaining children (n=14, Welch two-sample T-test: t(25.6) = -0.39, p = .70; Figure 4B top row, right handside). These results thus suggest lower myelination around left pOTS gaps, instead of the higher myelination predicted by hypothesis (i). This led to higher MD in the overall left pOTS in children with interruption - considering the cortex of the left pOTS proper and gyral gap(s) - than in children with continuous sulci (t(26.4) = -2.16, p = .040, Figure 4B bottom row). Additional analyses suggested that such differences were not driven by partial volume effects due to the presence of more or less surrounding white matter tissue or cerebro-spinal fluid close to gyral or sulcal cortex (see Supplementary DTI analyses in Appendix A).

The pattern of results observed for mean MD in the cortex was moreover confirmed by supplementary analyses of Radial Diffusivity (RD) and Fractional Anisotropy (FA) in the cortex around the left pOTS, and of MD, RD, and FA in the superficial white-matter located above the cortex (FA and RD being relevant markers of myelin content in white matter; see Dubois et al., 2014; Lazari & Lipp, 2021), suggesting lower myelination both in and above gyral gaps (Supplementary Figure 8 and Supplementary Table 3). In short, against hypothesis (i), the present data do not support the idea that left pOTS interruptions are associated with more terminations of long-distance projection fibers.

Next, we tested hypothesis (ii) according to which left pOTS gaps would be associated with increased local connectivity surrounding the left pOTS, through the packing of more U-shaped fibers traveling superficially. In order to compare the structural connectivity around the left pOTS proper and gaps in children with a continuous vs. an interrupted sulcus, we performed whole-brain probabilistic tractography from a superficial white-matter layer located under the left pOTS and gap cortex (see Figure 4C for a seed region in an example participant). Mean structural connectivity from these seeds from all children showed strong local connectivity to fusiform and inferior temporal gyri adjacent to the superficial white-matter seeds above the left pOTS, and extending to the middle temporal gyrus, parahippocampal cortex, and anterior OTS (Figure 4C middle panel), consistent with the presence of short, neighborhood association fibers traveling under the sulcal cortex (Dannhoff et al., 2023; Reveley et al., 2015). When comparing structural connectivity between children with interrupted and continuous left pOTS through permutation testing, children with sulcal interruption showed higher local connectivity, as indexed by density maps reflecting the number of reconstructed streamlines, in a cluster located around the posterior section of the left pOTS, after controlling for demographic variables (age, gender and handedness, p<0.01 threshold-free cluster enhancement -TFCE-uncorrected, Figure 4C bottom panel, leftmost map), suggesting a higher development of short-distance connections, in agreement with hypothesis (ii).

In order to assess whether this higher local connectivity in children with left pOTS gaps may also directly participate in their reading benefit, a second permutation analysis was run modeling the presence of left pOTS interruption, T3 TOWRE reading scores, and demographic covariates. In this analysis, no statistical difference was observed between children with and without interruptions (at p<0.01 TFCE), but connectivity and reading scores positively correlated in a white matter cluster that partially overlapped with the connectivity cluster from the previous analysis (Figure 4C bottom panel, middle and rightmost maps). Despite the reduced statistical power, this suggests that advantages in local structural connectivity around the posterior left pOTS in children with sulcal interruption may be associated with better reading scores and constitutes a candidate pathway subtending the behavioral reading advantage.

## Discussion

To the best of our knowledge, the current study provides empirical evidence for the first time supporting that the presence of a gyral gap of the left pOTS before, or at, the onset of literacy instruction robustly and uniquely predicts better future reading skills throughout the first three years of elementary school. Importantly, this effect was observed prospectively and was shown to accumulate over time. It was specific to the left posterior section of the OTS and to the presence of interruptions rather than any other anatomical feature of sulcus. This marker of individual sulcal anatomy proved to be a large and independent contributor to shaping children’s reading abilities, with predictive performance comparable to or greater than that of well-established familial and pre-literacy cognitive measures. Cognitive benefits seemed relatively specific to reading, although significant associations were also observed for language, math and general cognitive skills intercorrelated with reading. Increased local U-fiber connectivity around the left pOTS associated with the presence of a gyral gap appeared as a likely mechanism underlying the behavioral benefits of such gaps on reading development. Below, we discuss these results in the context of (i) robust and specific longitudinal associations between left pOTS interruptions and reading behavior, (ii) precision approaches to the study of the individual differences in cognition, (iii) developmental origins of the impact of left pOTS interruption on reading, and (iv) common processes by which cortical constraints may influence cultural acquisitions.

### Robust and relatively specific longitudinal associations between left pOTS interruptions and reading behavior

Previous findings demonstrated a cross-sectional association between left pOTS interruptions and reading ability in children (Borst et al., 2016) and adults (Cachia et al., 2018). Building on this work, the present findings showed a large positive contribution of left pOTS interruption to reading abilities in a population of beginning readers, and crucially, demonstrate its longitudinal validity: sulcal interruptions at the onset of literacy instruction in 5-6 year-olds predicted children’s future reading skills at age 8.

Importantly, left pOTS interruptions were robustly associated with all aspects of reading tested at 8 years, ranging from single-word and pseudo-word naming (in both accuracy and speed) to sentence reading fluency and silent passage comprehension, as well as even spelling skills. Interestingly, the association between sulcal anatomy and reading was largest for passage comprehension - the most demanding and advanced aspect of reading.

Additionally, brain-behavior associations also extended to the most basic aspects of reading, e.g., letter recognition and orthographic processing, as demonstrated by significant associations between left pOTS interruption and letter knowledge already at age 5, on top of associations with timed word and pseudo-word reading (Figure 3B). This *general* effect on reading skills thus extends beyond previous reports, established on reading measures mostly tapping into phonological decoding skills (Borst et al., 2016; Cachia et al., 2018). Moreover, left pOTS morphology may have an effect on reading development across alphabets throughout a wide range of orthographic depths, as we replicate in English prior results in French and Portuguese (Borst et al., 2016; Cachia et al., 2018) - two scripts much more transparent in terms of grapheme-to-phoneme conversion.

Our study further extends previous results by Borst, Cachia and colleagues (2016; 2018) to a population of children with above-average cognitive and reading scores, from relatively high-SES families, and enriched for family history of dyslexia (cf. Table 1) - although no child met criteria for dyslexia by T3. While our sample is not representative of the general population, thereby limiting the direct generalizability of our results, half of the adult sample from Cachia et al. (2018) came from low socio-economic and rural backgrounds (matched to a sample of illiterate adults), suggesting that the association between left pOTS interruption and reading skills is robust across socio-economic contexts. Since our reviewed preprint came out, the association between left pOTS interruption and reading was further replicated (Cachia et al., 2025; Yao et al., 2026), notably in a larger child sample along the full reading continuum and oversampling children with dyslexia (Yao et al., 2026). (Of note, the finding that pOTS morphology correlated in adults with reading fluency rather than accuracy (Cachia et al., 2025) may also explain one of two prior negative findings concerning the association between pOTS and reading (Del Mauro et al., 2022)). Thus, our results together with the extant literature overall underline a common effect of left pOTS interruption observed across the continuum of reading abilities, and possibly of cognitive abilities.

Interestingly, left pOTS interruption also showed associations with other cognitive skills: it was associated with performance IQ at age 5 (see Figure 3B and E), and with performance IQ, math and language skills more broadly at age 8 (Supplementary Figure 6). However, left pOTS interruption was most strongly associated and clustered with assessments of reading at age 8 (as well as spelling and expressive language). In addition, the prospective association between left pOTS interruption and reading skills was not merely driven by an association with general cognitive abilities at age 5 (Figure 3C and F) and no association with any other pre-literacy skill was found (Figure 3B and E). Math, reading, language and cognitive skills are intrinsically highly intercorrelated, as we also observed at age 8, and are known to mutually and positively influence each other over time (Ünal et al., 2023; Peng & Kievit, 2020; Ferrer et al., 2007; Huettig & Hulstijn, 2025; Dehaene et al., 2015). The weaker pOTS cognitive associations may therefore result from the mere combination of a specific pOTS influence reading and of associations between reading and other cognitive domains, although more research is needed to better understand these associations and how they unfold during development. Together, these findings highlight a relative degree of specificity of left pOTS influences to reading abilities, in line with the well-documented functional specificity of the left pOTS for orthographic processing and reading (e.g., Dehaene & Cohen, 2011; Gaillard et al., 2006; Pillet et al., 2024; Wilson et al., 2013; Woolnough & Tandon, 2024; Zemmoura et al., 2015; Zhan et al., 2023).

### Precision approaches to the study of individual differences in cognition

Results of the present study add to a growing body of literature indicating that inter-individual differences in the morphology of gyral gaps in association cortices are associated with various aspects of cognition such as numerical processing and reading skills (Borst et al., 2016; Cachia et al., 2018, 2021; Roell et al., 2021; Santacroce et al., 2024; Schwizer Ashkenazi et al., 2024; Tissier et al., 2018). The present and previous findings relating gyral gaps to individual differences in cognitive performance require manual definitions of gyral gaps in a relatively small sample of individuals, consistent with a “precision imaging” approach (Cachia et al., 2021; Gratton et al., 2022; Rosenberg & Finn, 2022). While brain-behavior relationships, typically studied through automatically-extracted estimates such as cortical thickness or surface area, have been found to have small effect sizes and to require large N approaches (Marek et al., 2022), macro-anatomical features of neuroanatomy may be expected to have a larger impact and require smaller samples. Moving forward, however, future studies can integrate these two approaches by building tools to precisely measure gyral gaps in an automated fashion, which would result in both a large N, as well as precisely identified gyral gaps, in each individual using deep learning (Lyu et al., 2021; Rivière et al., 2022). Such large-N approaches are needed to answer a number of outstanding questions, such as, in our case, whether there is an additional behavioral reading benefit to having two or more left pOTS interruptions - which were too infrequent in our sample -, or whether gyral gaps placed in specific locations along the left pOTS may be optimal to facilitate reading acquisition.

### Developmental origins of the impact of left pOTS interruption on reading

Developmentally, the OTS starts to appear around 25-30 weeks of gestation (Chi et al., 1977; Garel et al., 2001; Hansen et al., 1993). Although the relative developmental timeline between the appearance of a sulcus and that of its interruptions remains unclear, interruption(s) of the OTS are thus thought to be stable features of neuroanatomy determined early on. In our study, we observed high reliability of the pOTS interruption from age 5 (T1) to age 8 (T3), which is consistent with the longitudinal stability of interruptions of (i) the intraparietal sulcus and postcentral sulcus (Schwizer Ashkenazi et al., 2024), (ii) sulci in inferior frontal cortex (Tissier et al., 2018), and (iii) patterns of the anterior cingulate sulcus (Cachia et al., 2016) observed in late childhood (approximately 8 years onwards) to early adulthood. Of course, genetic and environmental factors also likely contribute to the brain-behavior relationship identified in the present study. For example, a recent report has reported a marginal association in the interruption of the left OTS among mother-child dyads (Fehlbaum et al., 2022), suggesting potential contributions of genetic factors or of the familial environment in shaping left OTS morphology. Further, gyrification is significantly impacted by the prenatal environment (Amiez et al., 2018) and gestational age at birth (Dubois et al., 2019), notably in the case of regions surrounding the left pOTS (Papini et al., 2020; Schmitt et al., 2023). Multiple (e.g., twin) vs. single pregnancy and intrauterine growth restriction may also influence such gyrification patterns (Amiez et al., 2018; Dubois et al., 2008), each of which can be considered in future studies.

Moreover, in the present study, effects of left pOTS interruptions on reading skills accumulated and grew, as shown by significant serial mediation effects from T1 to T2 to T3 (Figure 2D). As such, left pOTS interruptions seem to have a continued beneficial influence on developing reading skills throughout the first few elementary grades. Whether such benefits to reading development continue to accumulate beyond 8 or 9 years of age remains to be explored, but these benefits seem to at least be retained later on, as suggested by reports in older children and adults (Borst et al., 2016; Cachia et al., 2018, 2025; Yao et al., 2026).

### Local connectivity differences at gyral gaps: a computational advantage?

The present findings empirically support that, despite variability in the location of individual gaps along the pOTS, children with at least one gyral gap have increased local connectivity in a posterior cluster of the left pOTS, compared to those with a continuous pOTS. Such increased connectivity could have a computational advantage. For instance, previous work has highlighted that junctions in gyral ridges, known as 3-hinge gyri, such as those which surround gyral gaps, serve as hubs in cortico-cortical connections with higher connection degrees, stronger connections, and high betweenness centrality in graphs of structural connectivity (Zhang et al., 2020). The rich connectivity of such regions has also been documented through fine-grained post-mortem dissections (Shinohara et al., 2020; as also reviewed in Dannhoff et al., 2023), revealing that ‘*pyramid shape crossings*’ are characterized by, and seem to result from, the convergence of both intra- and inter-gyral U fibers. Crucially, we showed that such increased local U-shaped fibers also relate to reading skills.

Building on these results, mechanistically, computational simulations of white-matter development and gyrification suggest that gyral regions with a high concentration of growing axonal fibers tend to form 3-hinge gyri (Ge et al., 2018; see also Chavoshnejad et al., 2021). Therefore, pOTS gyral gaps may be intrinsically related to better U-fiber connectivity at the level where such gaps meet the adjacent fusiform and inferior temporal gyral crests. Such junctions may serve as connectivity hubs for processing written words, and such hubs may result in a computational advantage related to skilled reading, which can be tested in future research.

### Common cortical constraints on cultural learning

Interestingly, associations between sulcal interruptions and behavior have also been reported for symbolic, but not non-symbolic, number processing (Roell et al., 2021; Schwizer Ashkenazi et al., 2024), highlighting common processes by which cultural acquisitions, too recent on the scale of human evolution to rely on dedicated neural circuits (Dehaene & Cohen, 2007), may be boosted by the presence pre-existing gyral gaps. In the case of the left pOTS, various other aspects of early brain structure have been proposed to constrain the development of reading skills and of functional activations for reading in the VWFA and broader neural network involved in reading such as anatomical connectivity (Bouhali et al., 2014; Grotheer et al., 2021; Li et al., 2020; Moulton et al., 2019; Saygin et al., 2016) and cortical morphology. In terms of the latter, longitudinal studies following pre-reading children at familial risk of dyslexia showed for instance that the surface area of the left fusiform gyrus (Beelen et al., 2019) and patterns of gray-matter volume around the VWFA (Skeide et al., 2016) could be predictive of future dyslexia diagnosis.

## Conclusion

Altogether, reading is a cultural invention too evolutionarily recent to have shaped brain structure. Instead, learning to read tunes pre-existing ventral occipito-temporal territories for written language (Dehaene & Cohen, 2007; Dehaene-Lambertz et al., 2018; Kubota et al., 2023). Thus, stable properties of children’s macro-anatomical architecture, such as the presence of left pOTS gyral gaps and the associated increase in white-matter connectivity, may guide the neural plasticity during such a critical cultural learning experience, impacting learning outcomes. Our present findings support the importance of cortical constraints on literacy learning and link longitudinal measures of local cortical folding, white matter properties, and reading ability with a precision imaging approach for the first time.

## Materials and Methods

### Participants

51 children were followed longitudinally from kindergarten to third grade, as part of an NIH-funded project (K23HD054720) focusing on children’s reading development. Specifically, children were assessed at three time points: T1 at a median age of 5.5 years, T2 at a median age of 7.1 years, and T3 around 8.2 years. Neuropsychological assessments, demographic and family information were collected at all time points, while only T1 and T3 included MRI acquisitions (see Figure 1A). Eight children were excluded from the main analyses assessing longitudinal associations between OTS and MFS morphology and reading acquisition: 4 children due to drop-out (did not come back for T3 visit), 1 due to neuroanatomical findings (large bilateral cysts in the anterior temporal lobe) and 3 due to excessive motion in the MRI at either T1 (2 children) or T3 (1 child). The resulting main sample consisted of N=43 children (58% males; 88% right-handed; see Table 1 for specific demographics). Children came from high socio-economic backgrounds (e.g., median average parental years of formal education: 17 years, equivalent to a master’s degree) and had above-average cognitive abilities and reading skills, despite 42% of children having family history of dyslexia (e.g., a parent or sibling diagnosed with dyslexia, see Table 1). No child had received a diagnosis of dyslexia by T3, nor did any child qualify for such a diagnosis based on the in-depth neuropsychological assessments conducted in the study.

The Institutional Review Boards of Stanford University, where data were collected, and of the University of California San Francisco, where data were analyzed due to transition of the principal investigator FH, approved the present study. Informed assent and consent were obtained from children and their guardians, respectively. The dataset included in this study has resulted in several prior publications on other aspects of the data (Black et al., 2012; Hosseini et al., 2013; Myers et al., 2014; Vandermosten et al., 2020; Xia et al., 2021; Yamagata et al., 2016).

### Behavioral tests

Children were tested at T1, T2, and T3 with an extensive battery of standardized neuropsychological assessments concerning literacy and pre-literacy skills. Reading skills were assessed at all time points with the Sight Word Efficiency (SWE) and Phonemic Decoding Efficiency (PDE) subtests from the Test of Word Reading Efficiency (TOWRE first edition, Torgesen et al., 1999). In these tests, children had to correctly read aloud as many items as possible in 45 seconds from lists of increasing difficulty of high frequency words, often with irregular grapheme-phoneme mappings (sight words), and of pseudo-words (phonemic decoding). We used TOWRE scores as the primary measure of reading scores in the current study as this speeded reading score was most similar to speeded reading tests that had previously shown associations with the sulcation of the OTS (Borst et al., 2016; Cachia et al., 2018). At T1, letter knowledge was assessed through the Letter Identification subtest of the Woodcock Reading Mastery Test R/NU (WRMT-R/NU, Woodcock, 1998). At T2 and T3, reading skills were also assessed with the Word Identification and Word Attack subtests from WRMT-R/NU, which respectively consist in untimed reading of low frequency words and of pseudo-words in isolation, with increasing difficulty. Additionally, the WRMT Passage Comprehension subtest required children to appropriately complete a sentence or a short passage choosing a word among multiple options (pictures are included for a third of the easiest items). Reading Fluency and Spelling subtests from the Woodcock–Johnson III Tests of Achievement (Woodcock, McGrew, et al., 2001) were also administered. In the Reading Fluency subtest, participants had to read a series of sentences and indicate for each whether it was true or false, within a 3-min time constraint. In the spelling subtest, children had to write down and correctly spell dictated words of increasing difficulty.

Children’s cognitive and pre-literacy skills were also assessed, notably with the following tests: The Concept Formation subtest of the WJ-III Tests of Cognitive Abilities (Woodcock, Mather, et al., 2001) was used at T1 and T3 as an estimate of general cognitive abilities, and more specifically performance intelligence quotient (pIQ). This learning task captures fluid reasoning by measuring children’s ability to identify and state the rule for why some colored geometric figures are in a box whereas others are not (corrective feedback and reinforcement of correct answers are provided).

Receptive vocabulary was measured with the Peabody Picture Vocabulary Test (fourth edition, Dunn & Dunn, 2007). Blending and Elision subtests from the Comprehensive Test of Phonological Processing (CTOPP first edition, Wagner et al., 1999) were used to derive a composite measure of phonological awareness. Rapid Automatized Naming (RAN) abilities were measured by averaging scores on the object and color subtests (Wolf & Denckla, 2005), to provide an estimate of rapid naming abilities independent from letter or symbol reading. Many other cognitive, academic and linguistic abilities were further assessed at T3 and served for a supplementary analysis - please refer to Supplementary Table 2 for a full list of tests and their description.

Information regarding family history of dyslexia was obtained using the Adult Reading History Questionnaire (ARHQ, Lefly & Pennington, 2000), for both parents (except for one father), and considering possible diagnosis of older siblings. Several measures of dyslexia family history were derived: a binary dyslexia family history variable (at least one parent or sibling with a dyslexia diagnosis, vs. none), mother and father ARHQ scores, as well as averaged parental ARHQ scores weighted by either total general or educational time spent with the child by each caregiver.

### MRI Data acquisition

MRI data were acquired on a 3T GE Healthcare scanner at the Richard M. Lucas Center for Imaging at Stanford University using an eight-channel phased-array head coil. T_1_-weighted (T_1_w) images were acquired at T1 and T3 with a fast spoiled gradient echo sequence (repetition time: TR = 8.52 ms; echo time TE = 3.42 ms; inversion time TI = 400 ms; flip angle FA = 15 °; NEX = 1; 128 slices; thickness = 1.2 mm; field of view FOV = 22 cm; in-plane resolution = 256 × 256; voxel size = 0.859 × 0.859 × 1.2 mm).

Diffusion-weighted images (DWI) were collected with a high-angular resolution diffusion-imaging (HARDI) protocol at Time 3 with a single-shot spin-echo, echo-planar imaging sequence (46 axial 3mm-thick slices; repetition time TR = 5000 ms; echo time TE = 81.7 ms; in-plane resolution = 128 × 128; voxel size = 2.0 × 2.0 × 3.0 mm^3^; 150 directions with b = 2500 s/mm^2^; 6 volumes with b = 0 s/mm^2^, posterior-to-anterior phase encoding). DWI data were successfully collected in 35 participants. Among these, one was excluded due to a neurological finding, 2 due to acquisition issues related to the bounding box (missing parts of the brain) and 3 for excessive motion (see DWI processing), resulting in a final DWI sample at T3 of 29 children.

### T_1_-weighted data processing

T_1_-weighted images were processed in Freesurfer (Dale et al., 1999; Fischl et al., 1999) version 7.1.1, using the high-resolution sub-millimeter pipeline (Zaretskaya et al., 2018), in order to (1) segment gray and white matter, (2) reconstruct gray and white matter surfaces, and (3) dissociate sulcal from gyral components. Images were first processed cross-sectionally for each scan. When images were available at both Time 1 and Time 3 for a child (N=43), the longitudinal Freesurfer pipeline was used, which first creates an unbiased subject average across time points (“base”) and then restarts multiple processing steps at each time point separately from the base template in order to refine tissue segmentations on each image (respectively T1 and T3 longitudinally processed images, Reuter et al., 2012; see Figure 1A). The application of such longitudinal processing steps has been demonstrated to result in more accurate assessments of anatomy and evaluation of longitudinal change, as we also confirmed. Segmentations were inspected visually for quality of the acquisition and of the Freesurfer segmentation bilaterally in ventral occipito-temporal cortex susceptible to affect the identification of the OTS and MFS. This led to the rejection of 1 participant at T1 and 2 subjects at T3 due to excessive motion artifacts. An additional participant was excluded due to large bilateral arachnoid cysts in the anterior temporal lobe.

### Sulcal definitions & criteria

#### Definition of the OTS and MFS

The OTS and MFS were identified manually, for each participant at each time point (T1 and T3), and processing pipeline (Freesurfer cross-sectional vs. longitudinal). Sulci were labeled by FB and verified or corrected by KW. Sulcal labeling was conducted without awareness of the children’s reading scores, and relied on the identification and the anatomy of nearby sulci based on previous criteria from our group (Weiner, 2019; Weiner et al., 2014) and others (Cachia et al., 2018). Surrounding sulci considered when defining the OTS and MFS included the collateral sulcus (CoS) medially, the posterior transverse collateral sulcus (ptCoS) posteriorly, the anterior transverse CoS (atCoS) anteriorly, and the inferior and superior temporal sulci (ITS and STS) laterally. Additionally, the pre-occipital notch (see Figure 1B) was referenced as a useful landmark identifying the posterior extent of the OTS on the pial surface. Sulci were defined on Freesurfer’s inflated surface as components with negative curvature, taking into consideration folding patterns as seen in the volume or white and pial surfaces, and potentially exclusive of connected branches corresponding to other sulci.

The ptCoS was first identified as the sulcus extending from the CoS-proper (Petrides, 2018) and forming the posterior boundary of the fusiform gyrus.

The OTS was then defined as the long and deep sulcus separating the fusiform gyrus from the inferior temporal gyrus. As such, its posterior boundary was defined by a virtual line extending from the ptCoS to the pre-occipital notch (as best identified on the pial surface), which delineates the boundary between temporal and occipital cortex (Duvernoy, 1999; Ono et al., 1990; Petrides, 2018). Note that, as a consequence, our OTS definition is exclusive of occipital components and branches, considered as the inferior occipital sulcus, although we retained the classical denomination of the *occipito*-temporal sulcus. In cases when the OTS extended into the occipital cortex, the OTS-proper was thus interrupted at the level of a *pli de passage* most consistent with the virtual line between the ptCoS and pre-occipital notch. The pre-occipital notch was not considered as part of the OTS and excluded from OTS delineations in the frequent case where it was connected to the OTS. Just as the ptCoS can be considered as the posterior hook identifying the posterior extent of the fusiform gyrus, the atCoS can be considered as an anterior hook identifying the anterior extent of the fusiform gyrus (Duvernoy et al., 1999). Laterally, the identification of the ITS, and of the STS and its posterior branches (Huntgeburth & Petrides, 2012) were used to disambiguate OTS components in complex cases.

The MFS was defined as the longitudinal sulcus that divides the fusiform gyrus into lateral and medial components following guidelines by Weiner and colleagues (Weiner et al., 2014; as also reviewed in Weiner, 2019). One characteristic feature of the MFS, due to its location in between the CoS and OTS on the cortical surface, is that it forms a distinct ω shape on coronal slices (corresponding to the middle bump of the ω). Indeed, as a putative tertiary sulcus, the MFS is about half as deep as the surrounding OTS and CoS (Weiner et al., 2014). Additionally, the anterior tip of the MFS consistently aligns with the posterior tip of the hippocampus (Grill-Spector & Weiner, 2014).

#### Posterior / anterior OTS

A fixed boundary in MNI template space at coordinate Y=-40 was used to distinguish posterior sections of the OTS (corresponding to typical coordinates of the VTC) from anterior sections. Posterior and anterior volumes in MNI space were first transformed to Freesurfer average subject space (*fsaverage*), then projected onto the *fsaverage* surface, and finally projected onto each individual surface, to split the OTS into posterior and anterior sections (pOTS and aOTS respectively, see Figure 1B).

#### Definition of gyral gaps / plis de passage for T3 DWI analyses

In order to investigate white-matter properties of the sulcal interruptions that may be associated with a reading benefit, the gyral gaps, including gyral crowns, located between chunks of the left pOTS were delineated manually by outlining the nearest points between the sulcal components (as shown in Figure 4).

### Sulcal measurements

From the defined OTS, pOTS, aOTS, and MFS, seven anatomical features were extracted as possibly relevant features: (i) sulcal interruption, (ii) interruption distance between sulcal components, (iii) sulcal length, (iv) sulcal depth, (v) depth of the sulcal pit, (vi) sulcal thickness, and (vii) sulcal surface area. Interruption distance between sulcal components was considered as Cachia and colleagues (2018) observed an association between the size of left pOTS interruptions along the postero-anterior axis and reading skills. In the present work, interruption distance was measured as the Euclidean distance between two sulcal components, or the sum of distances in instances where there were three or more components of a sulcus. Specifically, sulcal interruptions were counted as such when sulcal components were separated by at least 1mm, or when the total interruption distance across all components of a sulcus was larger than 1mm, as very small interruptions may be attributable to noisy scans, partial volume effects or poor brain segmentations. Sulcal length was considered as a complementary feature of sulcal interruption and interruption distance, known to be behaviorally relevant (see e.g., Parker et al., 2023, for associations between MFS length and facial recognition abilities). Sulcal length was measured by custom software as the largest geodesic distance between any two points within a given sulcal component, or the sum of lengths of all sulcal components in cases of interrupted sulci. Sulcal depth was measured using the *calcSulc* toolbox (Madan, 2019), by extracting the median Euclidean distance of the 8% deepest vertices of a given sulcus (as identified by Freesurfer *sulc* values) to the enclosing brain surface. We chose to select the 8% deepest vertices, rather than the initially set number of vertices (100 in the toolbox), in order to prevent the biasing of sulcal depth across sulci of different sizes (notably the much smaller MFS). Results were unchanged when using a percentage of deepest vertices ranging from 4 to 12%. The depth of the sulcal pit was also considered as sulcal pits often constitute stable functional and behavioral landmarks (e.g., Natu et al., 2021).

### DWI processing

Each diffusion-weighted image was first denoised using the Marchenko-Pastur principal component analysis technique implemented in MRTrix3 (Veraart et al., 2016). Because no diffusion data with reverse phase encoding was available, distortions due to susceptibility artifacts were corrected using the *Synb0-DISCO* approach (Schilling et al., 2020). Accordingly, an undistorted b0 image was estimated using deep learning (3D U-nets) in order to match the geometry of structural T_1_w images and intensity contrasts from diffusion images. The distortions were then estimated and applied using the *topup* tool in FSL (Andersson et al., 2003). Both distortion and motion corrections were applied using the *eddy* function in FSL (Andersson & Sotiropoulos, 2016). Due to relatively high motion in this pediatric sample, *eddy* was run with conservative options consisting in 8 iterations with decreasing amounts of smoothing at 10, 6, 4, 2, 0, 0, 0, 0 mm full width at half maximum (FWHM), with additional outlier detection and replacement, and estimation of susceptibility-induced field changes by subject movement (Andersson et al., 2018). Volumes with more than 10% of outlier slices as identified by *eddy*, or with more than 1 mm of displacement relative to the previous volume, were excluded. Three participants with more than 10% of discarded volumes were excluded from further analyses, resulting in a final sample of 29 children with DWI data at Time 3.

In order to most accurately align individual T_1_w and DWI data, non-linear transformations were estimated using Advanced Normalization Tools (ANTs, Avants et al., 2011) between the participant’s bias-corrected T_1_w image and the individual average of all distortion-corrected b0 images (which have a relatively high gray/white contrast compared to other DWI-derived images). Specifically, non-linear transformations were used as an additional measure to correct for distortions induced in VTC by susceptibility artifacts due to the proximity of air/tissue interfaces in the nearby ear canals.

DWI data were first used to derive diffusion tensor indices in order to compare microstructural properties of the cortex and white matter around the left pOTS between sulcal components per se and the gyral gaps, and between children with and without interruptions. Second, probabilistic tractography from this region was performed to investigate the correlates of pOTS interruptions in terms of structural connectivity, between groups of children and according to reading scores.

The tensor model was estimated in FSL using *dtifit* in order to derive diffusion-tensor imaging (DTI) maps of mean and radial diffusivities (MD and RD) and fractional anisotropy (FA), as indices of myelin content. While MD is the most robust DTI index reflective of myelination and neurite properties in the cortex (Fukutomi et al., 2019), FA and RD are relevant markers of myelin content in white matter (Dubois et al., 2014; Lazari & Lipp, 2021).

To estimate MD, RD, and FA values most accurately in the cortical ribbon and superficial white matter above the left pOTS and its interruptions, individual MD, RD, and FA maps were transformed into native T_1_w space and averaged within local masks of the cortical ribbon and superficial white matter. These masks were obtained by propagating Freesurfer labels of the left pOTS and its interruptions into the volumes using the *mri_label2vol* function, either into the whole cortical ribbon or into a 2-voxel-thick layer of the superficial white matter (see Figure 4B and C). Additionally, a white matter mask was used to prevent the white-matter layer from extending into gray matter voxels of neighboring sulci.

To identify differences in structural connectivity associated with interruption of the left pOTS, probabilistic tractography was performed from a mask of superficial white matter located above the left pOTS labels, inclusive of the gyral crowns of the interruption(s) / *pli(s) de passage* in children with left pOTS interruption. Masks of superficial white matter above the left pOTS and interruptions were transformed from T1-w images into diffusion native spaces through ANTs using nearest-neighbor interpolation, to serve as tractography seeds. FSL’s *bedpostx* function (Bayesian Estimation of Diffusion Parameters Obtained using Sampling Technique) was used to estimate distributions of diffusion parameters and to model crossing fibers in each voxel (3 fibers per voxel). *Probtrackx* was used to perform probabilistic tractography from individual seeds, using the Euler integration for the precise computing of probabilistic streamlines, distance correction to compensate for the fact that connectivity distribution drops with distance from the seed mask, and with segmentations of gray matter and ventricles as termination masks. The minimum allowable angle between two tractography steps was set to 30 degrees, to allow for the reconstruction of U-shaped fibers, which pass by the white-matter underlying sulcal fundi connecting adjacent gyri - in this case, the fusiform and inferior temporal gyri. Because seed volumes significantly differed between children with interrupted and continuous left pOTS, the number of streamlines started per seed voxel in the tractography algorithm was scaled individually based on seed volume to correspond to an average of 5,000 seeds per voxel (default value in *probtrackx*) and to yield similar total numbers of streamlines. The latter was implemented in order to accurately assess group differences in structural connectivity, unbiased by seed volume. Individual masks of gray matter and ventricles were used as stop masks for the tractography. Distance correction and modified Euler streamlining options were used to improve the accuracy of tractography. The obtained individual probabilistic fiber density maps were then transformed into template MNI space for statistical analysis using non-linear registration in ANTs, from individual diffusion space (individual FA map) to the HCP1065 MNI FA template, and smoothed with a 4-mm-FWHM Gaussian kernel. Voxel-wise permutation (non-parametric) testing was performed with FSL’s *randomise* function (Winkler et al., 2014), to assess the effect of left pOTS interruption on structural connections of its underlying white-matter, controlling for age, gender and handedness. Results of such permutation tests were compared to those of a similar model with reading scores (controlled for demographic variables) as an additional predictor, so as to assess whether differences in structural connectivity were more attributable to interruption of the left pOTS or differences in reading skills.

### Statistical analyses

Statistical analyses were performed in R version 4.2.2 (R Core Team, 2022). The impact of sulcal anatomical features on reading skills, or other test scores, across time points was assessed while controlling for the possible impact of demographic variables (e.g., age, gender and handedness, on top of total brain volume (TBV) in the case of sulcal features). Specifically, for each sulcal feature and each assessment score of interest, an F-test was performed to compare two nested models predicting scores at time Ti (i=1, 2 or 3) from demographic variables only (baseline model), or taking into additional consideration sulcal features measured at time Tj (j= 1 or 3), controlled for nuisance variables, (test model) as follows:

Baseline model: Score_Ti_ ∼ Age_Ti_ + Gender + Handedness

Test model: Score_Ti_ ∼ Age_Ti_ + Gender + Handedness + residuals ( logit(sulcal interruption_Tj_ ∼ Age_Tj_ + TBV_Tj_ + Gender + Handedness))

where *logit* indicates a logistic regression.

Or, in the case of continuous sulcal measurements:

Test model: Score_Ti_ ∼ Age_Ti_ + Gender + Handedness + residuals(sulcal measure_Tj_ ∼ Age_Tj_ + TBV_Tj_ + Gender + Handedness)

F-statistics and associated p-values are reported for these nested model comparisons, as well as the difference in variance explained between models (ΔR^2^ in %). For more coherence with statistical tests, plotted scores are controlled for demographic variables (age, gender and handedness; i.e., residuals are plotted, after adding back population mean to retain the estimation of average scores, see e.g., Fig. 2a or Fig. S4). Plots were created using the R *ggplot2* package (Wickham, 2016). Early publicly available code corresponding to the R *ggrain* package was adjusted to create *raincloud plots* (Allen et al., 2019). To investigate the unique contributions of several predictors of reading (see for instance Figure S5b and f) while controlling for other variables, such predictors were added to both the baseline and test models, following the same logic. Finally, to investigate possible associations at T1 between sulcal interruption and precursors of reading, we compared the model fits between nested logistic regressions predicting left pOTS interruption based on demographic variables and TBV (baseline), including or not each reading precursor separately as follows:

Baseline model: logit(sulcal interruption_T1_ ∼ Age_T1_ + TBV_T1_ + Gender + Handedness)

Test model: logit(sulcal interruption_T1_ ∼ Precursor_score_T1_ + Age_T1_ + TBV_T1_ + Gender + Handedness)

Statistics were conducted on standard scores from the standardized neuropsychological tests as they are more normally distributed than raw scores (a requirement for parametric tests). More importantly, the rationale of the study is to assess whether sulcal morphology will predict whether a child is going to be a relatively good reader or not, independently from age. Note that despite standardization of raw scores within age bins in some tests, age still had a residual significant effect on test scores (TOWRE T3 standard score notably) and was therefore included in baseline models. Because the TOWRE is not normed below 6 years of age, an equivalent of a composite TOWRE standard score was obtained for T1 by scaling the principal component of T1 SWE and PDE raw scores (to a mean of 100 with standard deviation of 15).

To reduce degrees of freedom concerning measures of dyslexia family history, a stepwise backward selection was applied to identify family history measure(s) that best predicted T3 TOWRE composite scores (accounting for age, gender and handedness). Binary family history factor (i.e. *does a direct relative have a formal diagnosis?*), mother and father ARHQ scores and their weighted averages based on educational and total general parental time spent with the child were entered into the model. Among these, mom ARHQ scores were selected as the most reliable measure predicting children TOWRE T3 reading scores, without any significant additional contribution from other variables.

Because some children already knew how to read at Time 1 (median of 3 words read in 45s in the SWE subtest, ranging from 0 to 58; median of 2 syllables or pseudowords read in 45s in the PDE subtest, ranging from 0 to 36), complementary analyses were performed to (i) investigate whether the predictive power of left pOTS interruption held in a population of children with minimal reading skills at T1 (Fig. S4), and (ii) to better assess the effect size of independent contributions of left pOTS morphology on future reading skills (Fig. S5) over initial reading scores at T1 (to which it was already correlated). We therefore selected a subsample of children whose TOWRE standard score equivalent at Time 1 was below 95, resulting in a subset of 24 *initial minimal readers* (median of 0 words read in 45s in SWE, range: 0-7; median of 0 syllables in 45s in PDE, range: 0-4; see Table 1).

The serial mediation analysis was performed using the R *lavaan* package (Rosseel, 2012). Parameter estimates and confidence intervals were estimated using 10,000 bootstrap samples.

## Acknowledgements

Authors would like to thank Olivier Coulon for useful discussion, as well as Jacob Miller for sharing his script to extract sulcal length. Data collection was funded by the National Institutes of Health (NIH) under grant K23HD054720 to FH. FB was supported by NIH grant R01HD094834, the Institute of Convergence for Language, Communication and the Brain (ILCB, grants from France 2030 ANR-16-CONV-0002 and the Excellence Initiative of Aix-Marseille University A*MIDEX), and the French Foundation for Medical Research (Fondation pour la Recherche Médicale, FRM ARF202209015734). JD was supported by the Fondation Médisite (under the aegis of the Fondation de France, grant FdF-18-00092867) and the IdEx Université de Paris (ANR-18-IDEX-0001). FH was supported by the National Science Foundation (NRT-2152202, MCA-2120888), NIH (R01DC013064, U24AT011281, R01HD094834, R01HD096261), U.S. Department of Agriculture’s (USDA) / National Institute of Food and Agriculture (NIFA) (2022-11791) and the Oak Foundation (OCAY-19-215). KSW was supported by the National Science Foundation (CAREER 2042251) and the Brain and Behavior Research Foundation (NARSAD 30738).

