## Appendix A for "Unique longitudinal contributions of sulcal interruptions to reading acquisition in children"

### Supplementary Information for *Unique longitudinal contributions of sulcal interruption to reading acquisition in children*

Florence Bouhali, Jessica Dubois, Fumiko Hoeft *, Kevin Weiner *

| Sulcal measure in the left pOTS | T1-T3 correlation cross-sectional processing (r, 95% CI) | T1-T3 correlation longitudinal processing (r, 95% CI) |
|---|---|---|
| Cortical thickness | 0.636 [0.415, 0.786] | 0.742 [0.568, 0.786] |
| Surface area * | 0.936 [0.884, 0.965] | 0.985 [0.972, 0.992] |
| Sulcal length * | 0.659 [0.447, 0.801] | 0.977 [0.957, 0.987] |
| Total interruption distance * | 0.794 [0.648, 0.883] | 0.98 [0.963, 0.989] |
| Sulcal depth | 0.974 [0.951, 0.986] | 0.988 [0.978, 0.994] |
| Depth of sulcal pit | 0.934 [0.882, 0.964] | 0.975 [0.954, 0.987] |

**Supplementary Table 1: T1-T3 measurement stability of several sulcal measures in the left pOTS** (as an example)**, indexed by Pearson correlation coefficients across time points, based on Freesurfer cross-sectional vs. longitudinal processing.** (* denotes measures that show significantly higher T1-T3 reliability using longitudinal compared to cross-sectional processing, as shown by non-overlapping confidence intervals.)

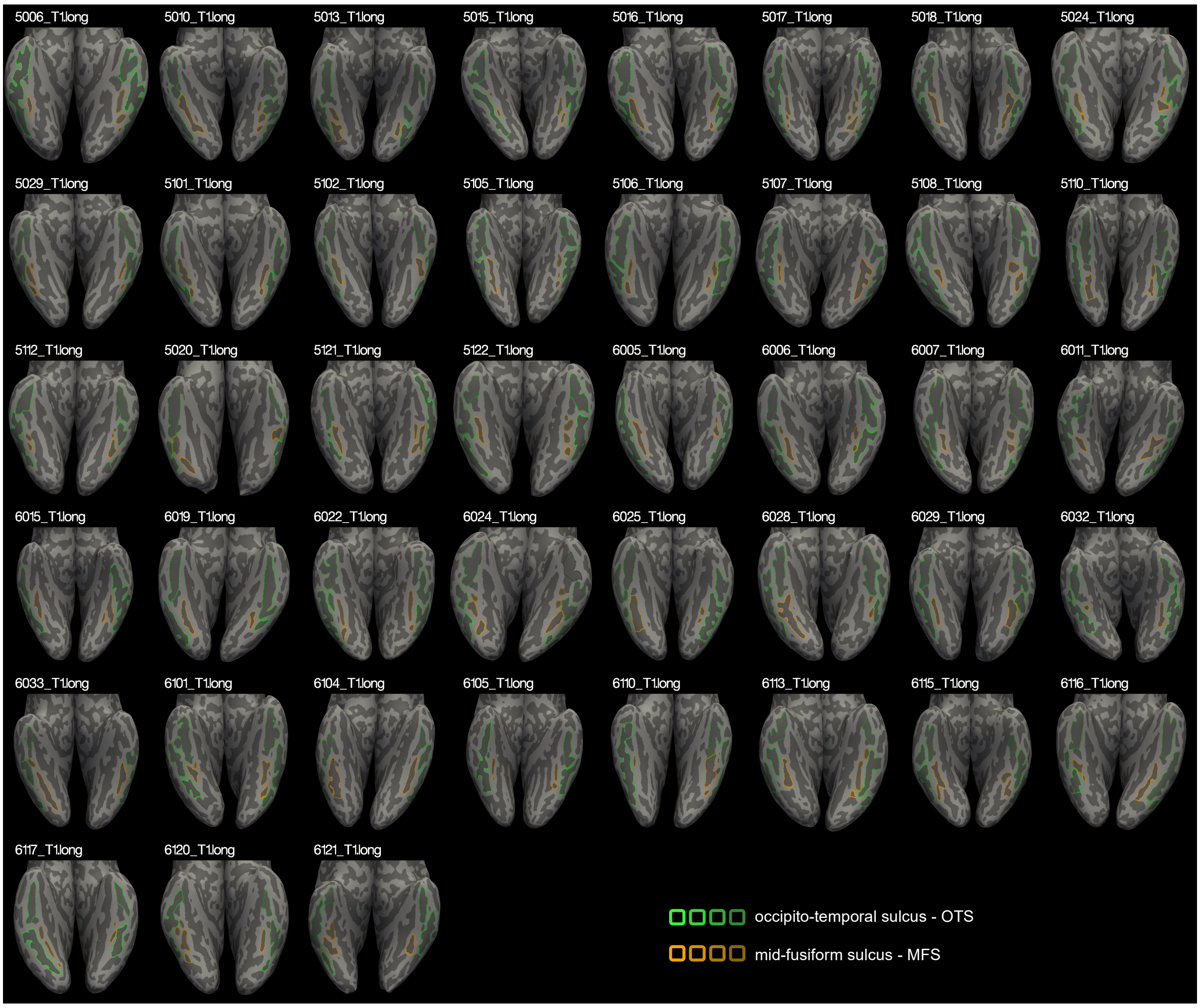


**Supplementary Figure 1: Sulcal outlines of the OTS and MFS in all 43 children with longitudinal MRI data at Time 1.**

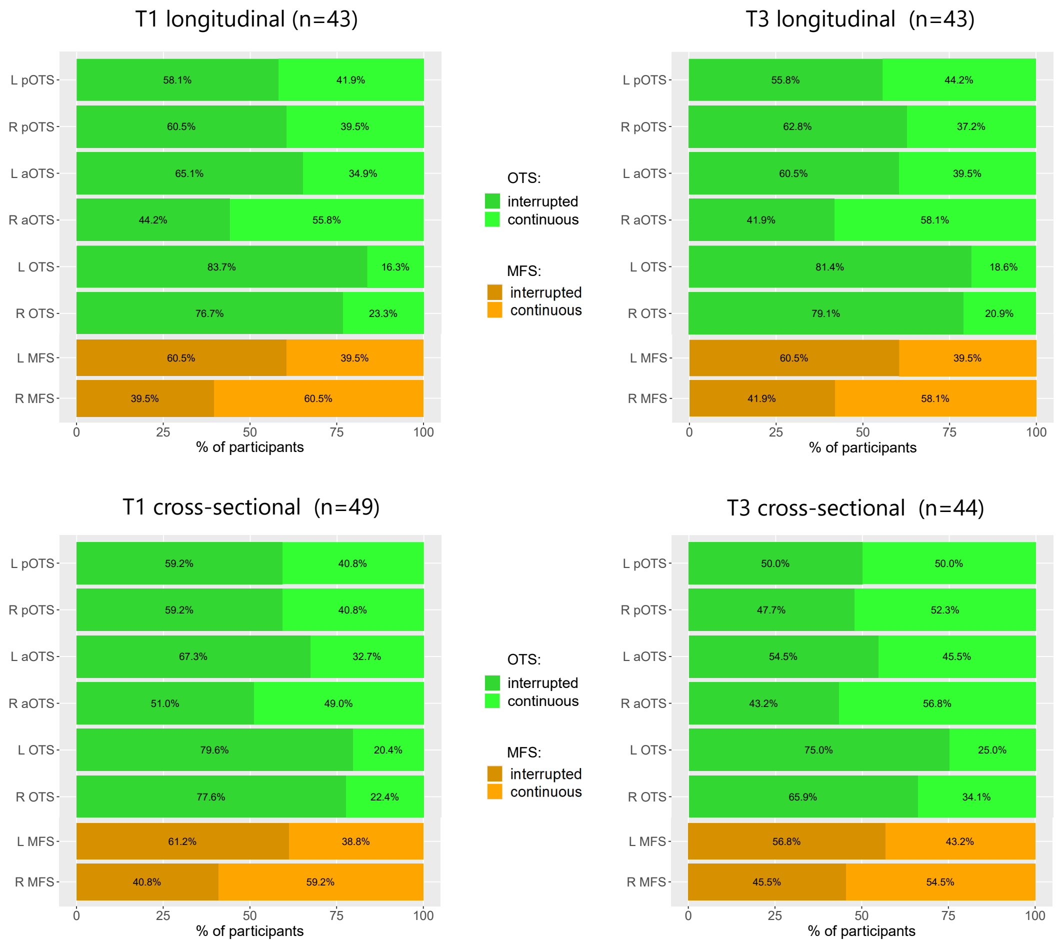


**Supplementary Figure 2: Incidence of interruptions of the OTS and the MFS at the different time points (T1 and T3), using both Freesurfer longitudinal and cross-sectional pipelines.** (OTS: full occipito-temporal sulcus; pOTS/aOTS: posterior/anterior sections of the OTS, relative to a boundary located at Y=-40 in MNI template space; MFS: mid-fusiform sulcus; L/R: left/right.)

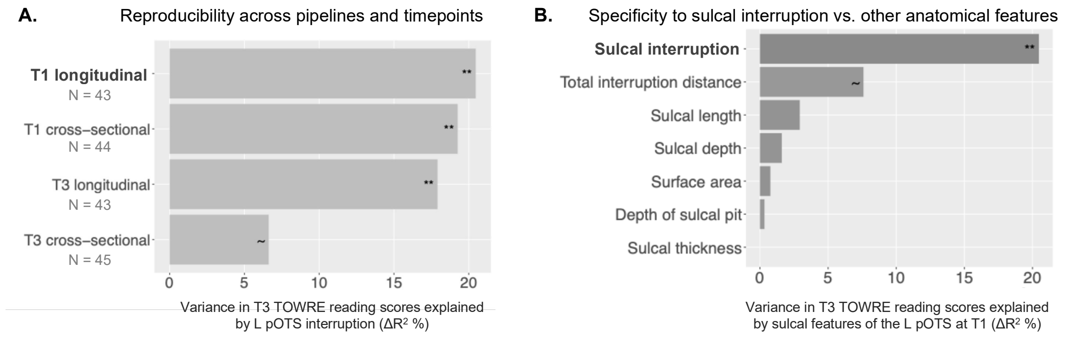


**Supplementary Figure 3: Reproducibility and specificity of the association between left pOTS interruption and reading scores. A.** Reproducibility across pipelines and time points: percentage of variance in TOWRE T3 reading scores explained by left pOTS interruption as measured at T1 or T3 through cross-sectional or longitudinal processing ($\Delta R^2$; respective N indicate the sample size for each analysis). **B.** Percentage of variance in TOWRE T3 reading scores explained by the different sulcal features of the left pOTS considered.

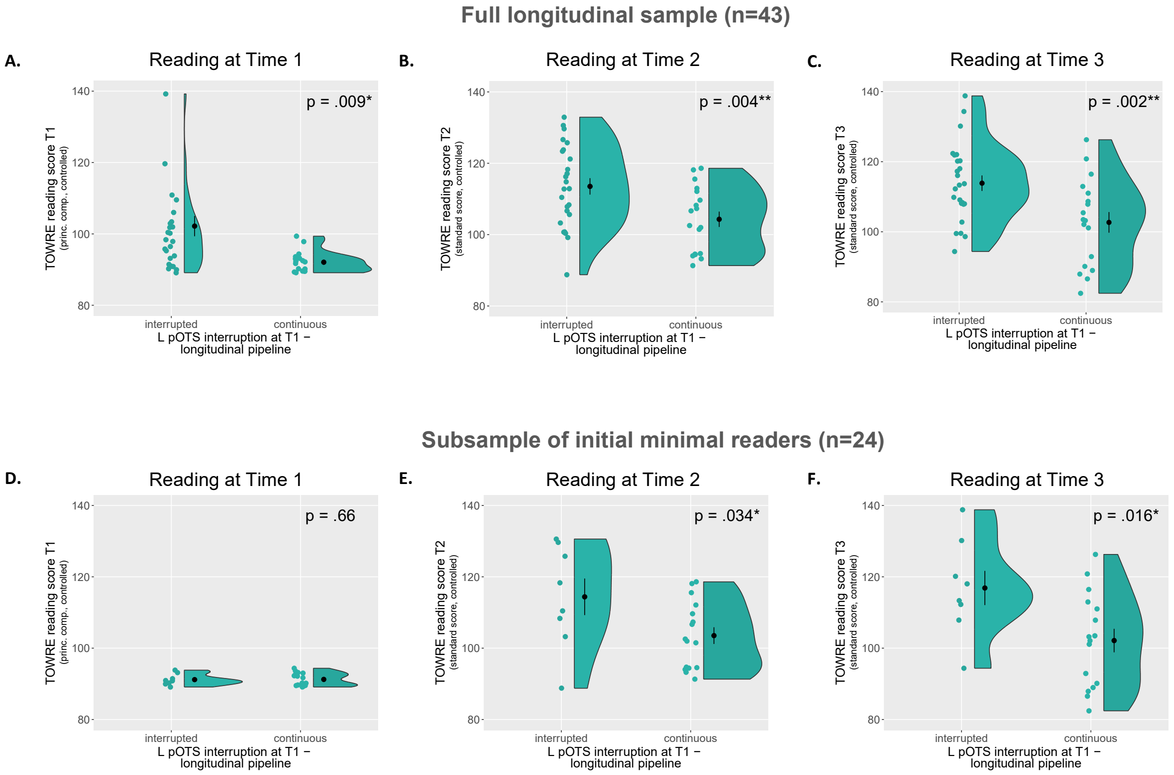


**Supplementary Figure 4: Differences in reading scores (TOWRE) at Times 1, 2 and 3 between children with interrupted or continuous left pOTS** (as measured at Time 1 with the longitudinal processing pipeline). **(A-C)**: Differences in reading for all children with longitudinal MRI data (N=43). (Panel c is duplicated from Figure 2A). **(D-F)**: Differences in reading scores in a subset of 24 children with lowest reading scores at Time 1, considered as initial minimal readers. Of note, despite the absence of a difference in reading scores at Time 1 between children with continuous and interrupted left pOTS (d), children with interrupted left pOTS read significantly better at Time 2 (e) and Time 3 (f) than those with continuous left pOTS in this sample.

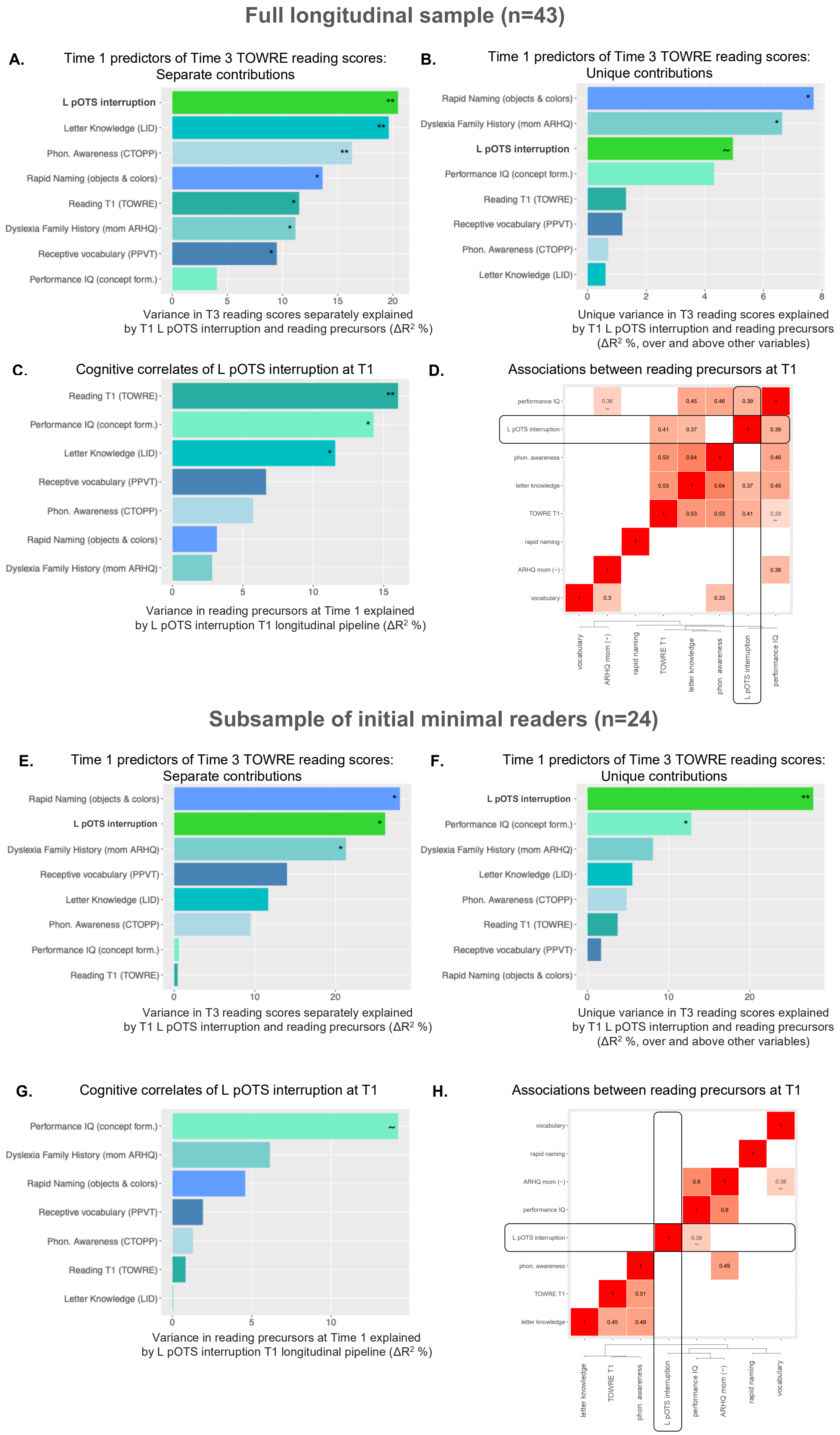

Full longitudinal sample (n=43)
A.
Time 1 predictors of Time 3 TOWRE reading scores: Separate contributions
L pOTS interruption
Letter Knowledge (LID)
Phon. Awareness (CTOPP)
Rapid Naming (objects & colors)
Reading T1 (TOWRE)
Dyslexia Family History (mom ARHQ)
Receptive vocabulary (PPVT)
Performance IQ (concept form.)
Variance in T3 reading scores separately explained by T1 L pOTS interruption and reading precursors (ΔR² %)
B.
Time 1 predictors of Time 3 TOWRE reading scores: Unique contributions
Rapid Naming (objects & colors)
Dyslexia Family History (mom ARHQ)
L pOTS interruption
Performance IQ (concept form.)
Reading T1 (TOWRE)
Receptive vocabulary (PPVT)
Phon. Awareness (CTOPP)
Letter Knowledge (LID)
Unique variance in T3 reading scores explained by T1 L pOTS interruption and reading precursors (ΔR² %, over and above other variables)
C.
Cognitive correlates of L pOTS interruption at T1
Reading T1 (TOWRE)
Performance IQ (concept form.)
Letter Knowledge (LID)
Receptive vocabulary (PPVT)
Phon. Awareness (CTOPP)
Rapid Naming (objects & colors)
Dyslexia Family History (mom ARHQ)
Variance in reading precursors at Time 1 explained by L pOTS interruption T1 longitudinal pipeline (ΔR² %)
D.
Associations between reading precursors at T1
performance IQ
L pOTS interruption
phon. awareness
letter knowledge
TOWRE T1
rapid naming
ARHQ mom (−)
vocabulary
Subsample of initial minimal readers (n=24)
E.
Time 1 predictors of Time 3 TOWRE reading scores: Separate contributions
Rapid Naming (objects & colors)
L pOTS interruption
Dyslexia Family History (mom ARHQ)
Receptive vocabulary (PPVT)
Letter Knowledge (LID)
Phon. Awareness (CTOPP)
Performance IQ (concept form.)
Reading T1 (TOWRE)
Variance in T3 reading scores separately explained by T1 L pOTS interruption and reading precursors (ΔR² %)
F.
Time 1 predictors of Time 3 TOWRE reading scores: Unique contributions
L pOTS interruption
Performance IQ (concept form.)
Dyslexia Family History (mom ARHQ)
Letter Knowledge (LID)
Phon. Awareness (CTOPP)
Reading T1 (TOWRE)
Receptive vocabulary (PPVT)
Rapid Naming (objects & colors)
Unique variance in T3 reading scores explained by T1 L pOTS interruption and reading precursors (ΔR² %, over and above other variables)
G.
Cognitive correlates of L pOTS interruption at T1
Performance IQ (concept form.)
Dyslexia Family History (mom ARHQ)
Rapid Naming (objects & colors)
Receptive vocabulary (PPVT)
Phon. Awareness (CTOPP)
Reading T1 (TOWRE)
Letter Knowledge (LID)
Variance in reading precursors at Time 1 explained by L pOTS interruption T1 longitudinal pipeline (ΔR² %)
H.
Associations between reading precursors at T1
vocabulary
rapid naming
ARHQ mom (−)
performance IQ
L pOTS interruption
phon. awareness
TOWRE T1
letter knowledge

**Supplementary Figure 5: Longitudinal contributions of left pOTS interruption to reading relative to cognitive precursors of reading, and their associations,** in the full longitudinal sample (A-D, top) and in the sub-sample of initial minimal readers (E-H, bottom), in which left pOTS interruption and T1 reading skills are unconfounded. **A & E.** Variance in TOWRE T3 reading scores explained by left pOTS interruption and classical predictors of reading at Time 1 separately, over and above demographic variables, ranked by decreasing contributions ($\Delta R^2$ in percentage, *a* is duplicated from Figure 3a). Baseline reference models with only demographic variables account respectively for a total $R^2$ of 11.0 and 2.2% of variance in T3 reading. **B & F.** Unique variance in TOWRE T3 reading scores explained by left pOTS interruption and classical predictors of reading at Time 1, over and above other variables of interest and demographic variables, ranked by decreasing contributions ($\Delta R^2$ in percentage). Reference models with all considered variables account respectively for a total $R^2$ of 58.9 and 75.8%of variance in T3 reading. **C & G.** Associations between left pOTS interruption and predictors of reading at Time 1: percentage of variance ($\Delta R^2$) in left pOTS interruption explained by each pre-literacy skill separately, over and above demographic variables and total brain volume (panel c is duplicated from Figure 3b). **D & H.** Correlation matrices showing significant associations between precursors of reading at T1, and their associated clustering dendrogram shown on the x-axis. (Mother ARHQ scores were multiplied by -1 so that higher values would be associated with better cognitive skills, in keeping with other variables. p-values for associations located above the diagonal are corrected for multiple comparisons at a False Discovery Rate. Panels A-C and E-G are reproduced from Figure 3 to facilitate comparison. (**: p<0.01, *: p<.05, ~:p<.07)

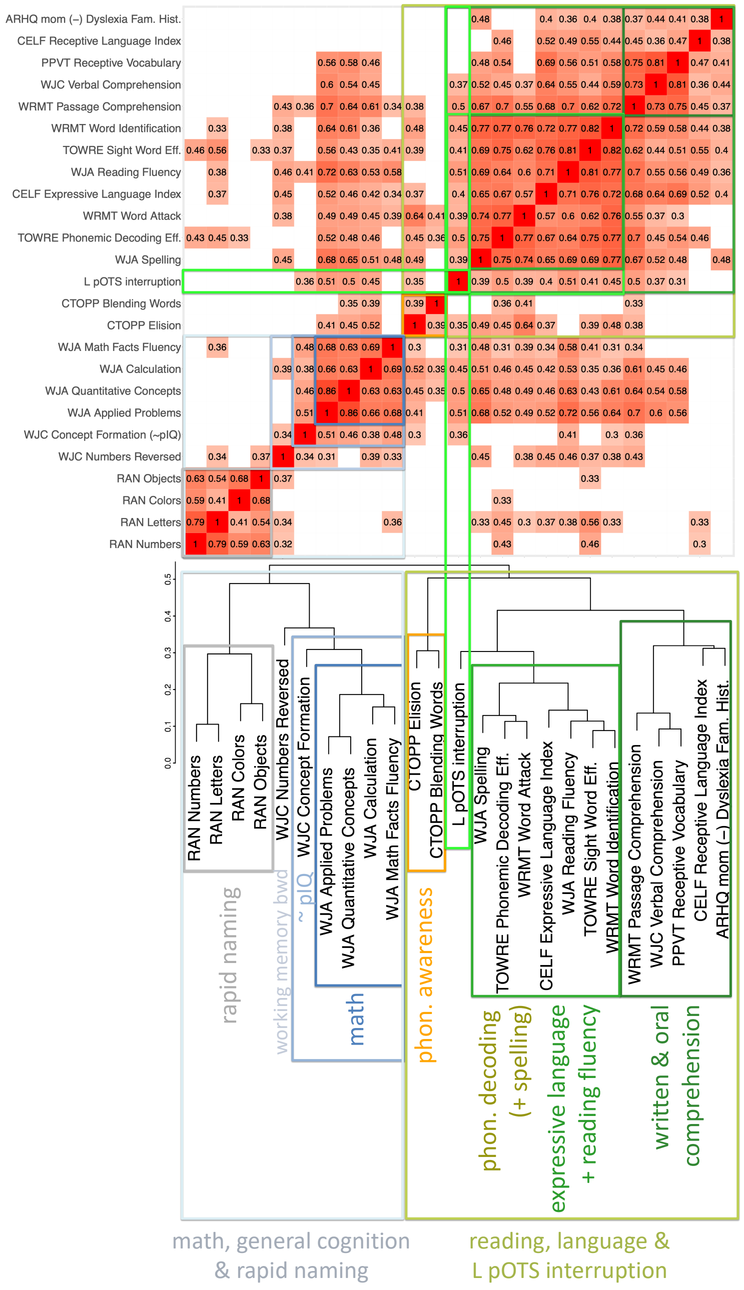


**Supplementary Figure 6: The analysis of the cognitive correlates of left pOTS interruption at Time 3 shows a relative cognitive specificity of this sulcal marker to reading.**
Correlation matrix showing the associations between cognitive and linguistic assessments at T3 and left pOTS interruption. The correlation matrix was ordered by hierarchical clustering with complete linkage, using a Pearson correlation-based distance matrix defined as (1 – r) / 2. The associated clustering dendrogram is displayed below the matrix x-axis, with the different clusters highlighted in the same color in the matrix and dendrogram.
Only significant correlations are shown (p<.05), with uncorrected p-values below the diagonal and FDR-corrected p-values above the diagonal. (The residuals of the logistic regression of left pOTS interruption by demographic variables and total brain volume were used for this analysis, as in previous analyses.)

**Supplementary Table 2: Description of the neuropsychological assessments included in the analysis of the cognitive correlates of left pOTS interruption at Time 3**, listed by battery. Standardized scores were included for each subtest, except for the CELF assessments of oral language for which receptive and expressive language composite scores were used due to numerous subtests.

| | |
|---|---|
| **Woodcock–Johnson III Tests of Achievement** | |
| WJA Reading Fluency | Timed silent reading of simple sentences. Decide quickly if sentences are true/false. Measures speed and accuracy. |
| WJA Spelling | Write increasingly difficult words. Measures ability to spell correctly. |
| WJA Calculation | Perform additions, subtractions, multiplications and divisions of increasing difficulty. Measures accuracy and understanding of mathematical procedures and concepts. |
| WJA Math Facts Fluency | 3 min to complete as many short arithmetic problems as possible. Measures automaticity and fluency in recalling basic math facts. |
| WJA Applied Problems | Listen to problems, identify key information and procedure, then perform simple calculations. Measures ability to analyze and solve math problems. |
| WJA Quantitative Concepts | Oral questions assessing understanding of mathematical concepts, such as number knowledge, mathematical vocabulary, and recognition of mathematical symbols and operations. Measures comprehension of foundational mathematical ideas rather than computational skills. |
| **Woodcock–Johnson III Tests of Cognitive Abilities** | |
| WJC Concept Formation | Controlled learning task which measures the ability to absorb and apply learning from basic principles. Identify and state the rule for a concept about a set of colored geometric figures when shown instances and non-instances of the concept. |
| WJC Visual Matching (2) | 3 min to locate and mark matching pairs of numbers within rows of 6 numbers (increasing difficulty from single- to triple-digit numbers). Measures visual processing speed. |
| WJC Verbal Comprehension | Measures knowledge of word meanings. Combines naming pictured objects, providing synonyms or antonyms, and completing sentences to indicate an appropriate analogy. |
| WJC Numbers Reversed | Repeat recorded series of numbers backward. Measures short-term auditory working memory. |
| **Peabody Picture Vocabulary Test 4** | |
| PPVT Receptive Vocabulary | The examiner says a word and the child points to the matching picture from a set of four. Measures receptive vocabulary / single-word spoken language comprehension. |
| **Rapid Automatized Naming RAN/RAS** | |
| RAN Letters | Name aloud a series of repeated single letters presented on a card as quickly and accurately as possible. Measures speed of automatic retrieval for letter names. |
| RAN Numbers | Name aloud a series of repeated single-digit numbers presented on a card as quickly and accurately as possible. Measures speed of automatic retrieval for number names. |
| RAN Colors | Name aloud a series of colored patches presented on a card as quickly and accurately as possible. Measures speed of automatic retrieval for color names. |
| RAN Objects | Name aloud a series of common objects presented on a card as quickly and accurately as possible. Measures speed of automatic retrieval for object names. |
| **Comprehensive Test of Phonological Processing** | |
| CTOPP Elision | The child hears a word and must say it with part removed (e.g., say baseball without ball). |
| CTOPP Blending Words | The child hears separate sounds and must blend them into a word (e.g., say /c/ /a/ /t/). |
| **Clinical Evaluation of Language Fundamentals** | |
| CELF Receptive Language Index | Composite score which measures spoken language comprehension in meaning and grammar. Subtests include: Sentence Structure (choose a picture that matches a spoken sentence) ; Concepts & Following Directions (listen to spoken directions of increasing length and complexity, then point to / identify items in the correct order); Word Classes Receptive (choose words that go together based on meaning). |
| CELF Expressive Language Index | Composite score which measures the ability to express ideas (grammar and vocabulary). Combines: Word Structure (complete sentences using correct grammatical forms - plurals, verb tense, pronouns, possessives); Formulated Sentences (create a complete, grammatically correct sentence using a given word); Recalling Sentences (auditory memory for language); Word Classes Expressive (explain why words go together based on meaning). |
| **Woodcock Reading Mastery Tests R/NU** | |
| WRMT Word Identification | Read aloud increasingly difficult real words presented in isolation. Measures sight-word reading accuracy and recognition of familiar printed words. |

| | |
|---|---|
| WRMT Word Attack | Read aloud increasingly difficult pronounceable pseudo-words. Measures knowledge of grapheme-phoneme correspondences and ability to apply phonics skills to decode new words. |
| WRMT Passage Comprehension | Read short passages or sentences and supply a missing word using contextual, vocabulary, and comprehension cues. Measures understanding of connected text and ability to derive meaning from what is read. |
| Test of Word Reading Efficiency - 1st edition | |
| TOWRE Sight Word Efficiency | Correctly read aloud as many words as possible in 45 sec from a list of high-frequency and often irregular words of increasing difficulty. |
| TOWRE Phonemic Decoding Eff. | Correctly read aloud as many pseudo-words as possible in 45 sec from a list of increasing difficulty. |
| Adult Reading History Questionnaire | |
| ARHQ mom (-) | Mother's score on a questionnaire about her reading history, assessing the mother's risk of dyslexia. The score is sign flipped so that higher score is associated with better performance, for consistency with other assessments. |

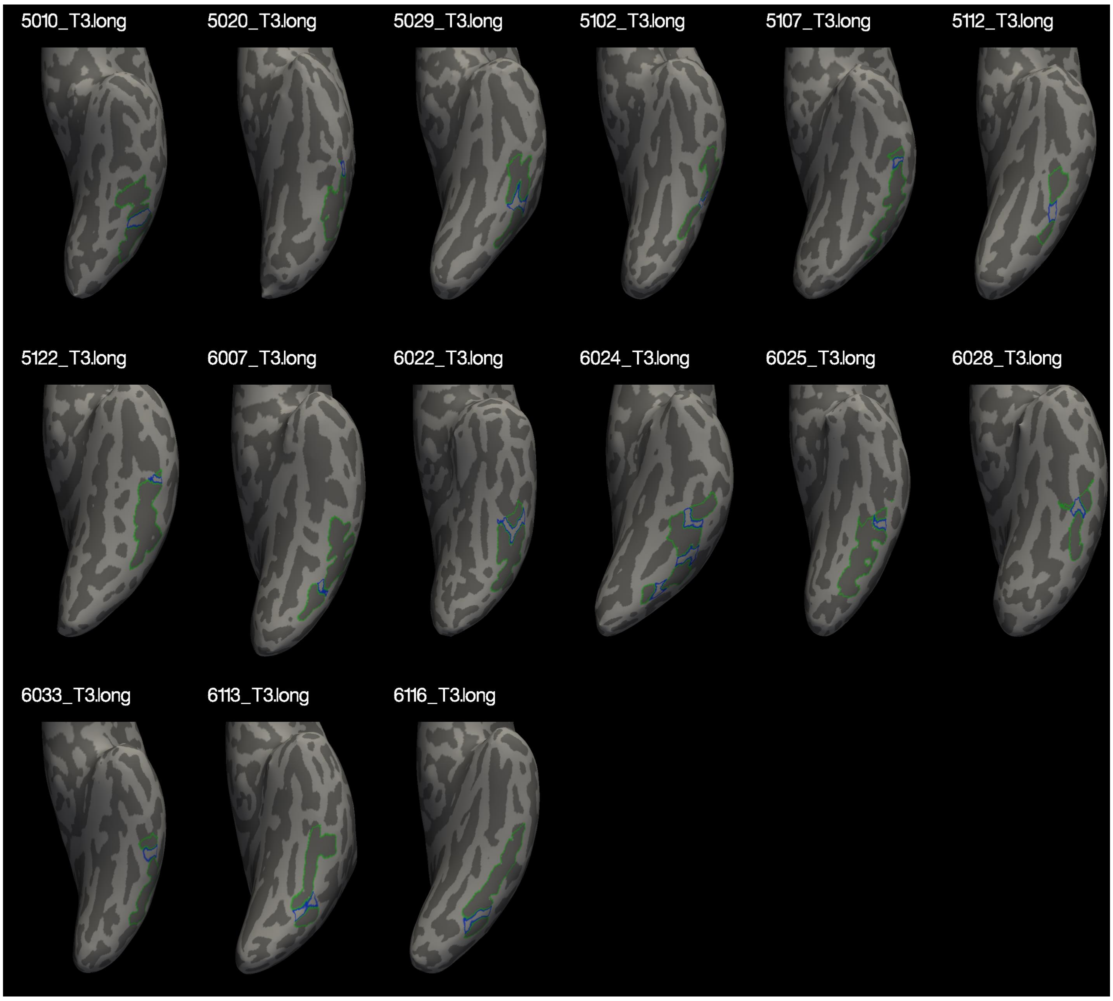


**Supplementary Figure 7: Sulcal outlines of the left pOTS** (green) **and left pOTS gyral gaps** (blue) in children with sulcal interruption(s) and diffusion data at Time 3 (Time 3 longitudinal processing).

#### Supplementary DTI analyses: controlling for partial volume effects

Despite the methodological care put in aligning diffusion maps to individual anatomy, observed differences in cortical MD between gyri and sulci could be artefactual, due to partial volume effects originating from the neighboring cerebro-spinal fluid (CSF) and superficial white matter. Indeed, CSF, which has high MD values, is present in larger quantities near the gyral than sulcal cortex, while sulcal cortex has more contact with underlying white matter characterized by lower MD values. Two supplementary analyses were therefore run to verify that our results were not driven by partial volume effects. (1) Median cortical MD values, instead of mean cortical MD, were derived individually throughout cortical voxels of the regions of interest, as medians would be less affected by voxels with outlier MD values. (2) Mean MD values were extracted from the middle half of the cortical ribbon (corresponding to cortical depths between 25 and 75%), to avoid contamination by CSF or white matter. Results of both control analyses yielded qualitatively similar results to that of mean MD throughout the cortical ribbon.

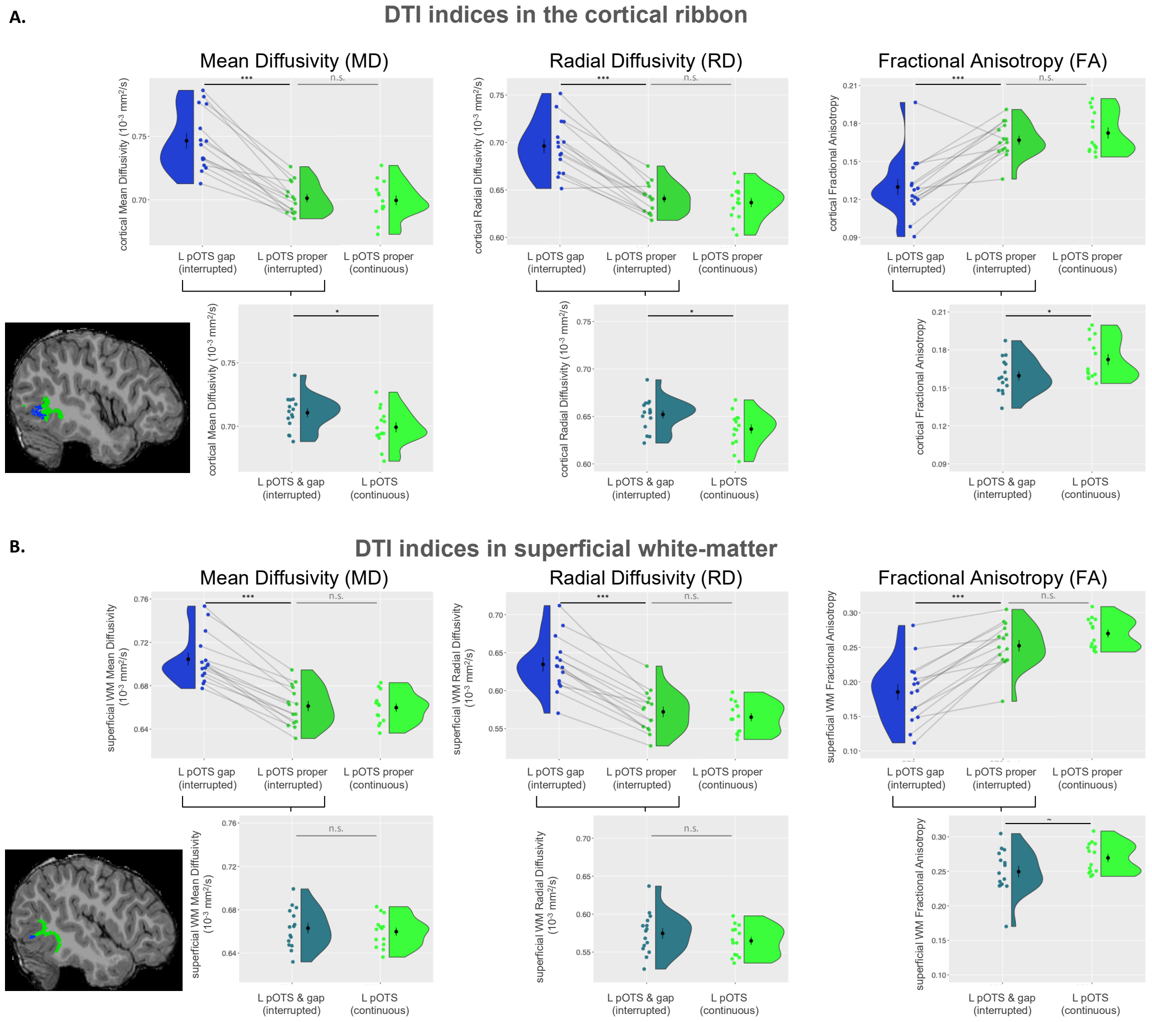


**Supplementary Figure 8: Diffusion-tensor properties around the left pOTS gaps and left pOTS proper** in children with an interrupted vs. continuous sulcus, in the cortical ribbon (A) and superficial white-matter (B).

| | L pOTS gap vs. proper (n=15 children with interruption, paired two-sample t-test: t(14), p-value) | | L pOTS proper: interrupted vs. continuous (n=15 and 14 children, Welch two-sample T-test: t, p-value) | | L pOTS (inc. gap if present): interrupted vs. continuous (n=15 and 14 children, Welch two-sample T-test: t, p-value) | |
|---|---|---|---|---|---|---|
| | Cortex | Superficial WM | Cortex | Superficial WM | Cortex | Superficial WM |
| MD | **9.74, $1.3*10^{-7}$** | **12.64, $4.8*10^{-9}$** | -0.39, .70 | -0.23, .81 | **-2.16, .040** | -0.52, .60 |
| RD | **9.49, $1.8*10^{-7}$** | **11.53, $1.6*10^{-8}$** | -0.64, .53 | -0.82, .41 | **-2.32, .028** | -1.11, .28 |
| FA | **-5.92, $3.7*10^{-5}$** | **-7.89, $1.6*10^{-6}$** | 1.01, .32 | 1.68, .11 | **2.19, .038** | 1.95, .062 |

**Supplementary Table 3: Statistical comparisons in DTI indices in the left pOTS and left pOTS gaps.** (MD: Mean Diffusivity, RD: Radial Diffusivity, FA: Fractional Anisotropy, WM: white matter; bolded text indicates significant differences.)
